# Structure of puromycin-sensitive aminopeptidase and polyglutamine binding

**DOI:** 10.1101/2023.05.30.542994

**Authors:** Sowmya Madabushi, K. Martin Chow, Eun Suk Song, Anwesha Goswami, Louis B. Hersh, David W. Rodgers

## Abstract

Puromycin-sensitive aminopeptidase (E.C. 3.4.11.14, UniProt P55786), a zinc metallopeptidase belonging to the M1 family, degrades a number of bioactive peptides as well as peptides released from the proteasome, including polyglutamine. We report the crystal structure of PSA at 2.3 Ǻ. Overall, the enzyme adopts a V-shaped architecture with four domains characteristic of the M1 family aminopeptidases, but it is in a less compact conformation compared to most M1 enzymes of known structure. A microtubule binding sequence is present in a C-terminal HEAT repeat domain of the enzyme in a position where it might serve to mediate interaction with tubulin. In the catalytic metallopeptidase domain, an elongated active site groove lined with aromatic and hydrophobic residues and a large S1 subsite may play a role in broad substrate recognition. The structure with bound polyglutamine shows a possible interacting mode of this peptide, which is supported by mutation.

## Introduction

Bioactive peptides perform a variety of signaling functions [1, 2]. As neuromodulators, they play a vital role in regulating activity throughout the nervous system, while in the periphery they maintain normal function in organ systems as well as modulate responses to environmental stimuli. A number of peptidases have been implicated in the control of bioactive peptide levels [1–4], and nearly all of these neuropeptidases utilize a zinc ion cofactor with catalytic domains structurally related to the well-characterized enzyme thermolysin [5]. A group of these peptidases belong to the MA clan, which is characterized by a HEXXHX_18_E active site sequence motif, where the two histidines and the distal glutamate coordinate the zinc ion cofactor. A water molecule that also coordinates the zinc acts as the attacking nucleophile in catalysis. It is stabilized by hydrogen bonding to the first glutamate of the motif, which also serves as a general base to abstract a proton from the water, facilitating nucleophilic attack.

Within clan MA, members of the M1 family are aminopeptidases characterized by a second conserved sequence, GAMENW, in which the glutamate residue has been proposed to interact with the amino terminus of substrates [6]. M1 member puromycin-sensitive aminopeptidase (PSA or NPEPPS, E.C. 3.4.11.14, UniProt P55786), named for its inhibition by the antibiotic puromycin, accounts for the major fraction of cytosolic aminopeptidase activity in various human tissues [7]. PSA was first identified based on its metabolism of opioid peptides [8–10], and it has been implicated in cell cycle control [11, 12] development of cell polarity [13–16], and processing of peptides released by the proteosome [17] as well as being identified as a target for cancer therapies [18, 19].

PSA deficient mice, known as Goku mice, have been generated by the gene trap method [20]. Both male and female Goku mice exhibit reproductive defects. The male Goku mice are infertile and lack copulatory behavior. Female Goku mice also show infertility due to impaired formation of the corpus luteum during pregnancy [21]. Deletion of the PSA orthologs in *C. elegans* and *Drosophila* also produces effects on reproduction and embryonic development [14, 22–24]. The Goku mice in addition exhibit compromised pain perception and increased anxiety, which may result from changes in the levels of circulating enkephalins [25].

More recently, PSA gene expression in various regions of mouse brain was shown to correlate with low levels of expressed mutant human Tau protein, and overexpression of PSA inhibited Tau induced neurodegeneration in a *Drosophila* model [26]. A direct role for PSA in metabolizing Tau has been proposed [26–29], although PSA cannot degrade Tau *in vitro* [30]. Thus, its precise role in Tau regulation remains unknown. PSA performs the important function of degrading polyglutamine peptides released by proteasomes [31],and loss of PSA activity results in increased aggregation and toxicity of polyglutamine expanded Huntingtin exon 1 in cultured cells and muscle [32]. Loss of PSA also affects SOD1 abundance and clearance, suggesting a neuroprotective role in amyotrophic lateral sclerosis [33]. PSA is therefore particularly associated with neurodegenerative disorders as well as normal peptide metabolism and reproductive function.

We present here a crystal structure of PSA that defines the architecture of the enzyme and the mechanism underlying its restriction to exopeptidase activity. The structure also suggests a basis for the enzyme’s ability to degrade a broad range of peptide sequences. In addition, we define the binding path of a polyglutamine peptide in the active site of the enzyme.

## Materials and methods

### Production of PSA

Human PSA for crystallization was expressed in insect cells using the BAC-TO-BAC system (Invitrogen) as described previously [34, 35]. The coding sequence for PSA was introduced into the pFASTBAC-HT(B) intermediate vector, which codes for a polyhistidine affinity tag and a TEV protease site on the N-terminus of the protein. Protein was produced by culturing suspended Sf9 insect cells in sf-900 II serum free medium (Gibco BRL) at 27°C.

Expressed PSA was purified by anion exchange chromatography using POROS HQ resin (GE Healthcare) with the sample and resin initially equilibrated with 20 mM Tris (pH 7.4). The enzyme was eluted with a gradient of increasing NaCl concentration ranging from 0.1 M to 1 M. Peak fractions corresponding to PSA were concentrated and run through a molecular sieving column (Sephadex G50) equilibrated with 20 mM Tris (pH 7.4). Fractions containing PSA were pooled, dialyzed against 10 mM HEPES buffer (pH 7.0), 2.0 mM BME and concentrated to 5-8 mg/ml of apparently homogenous enzyme. The N-terminal sequence was not removed for crystallization trials. Before obtaining kinetic data for PSA, we switched to producing the enzyme in insect cells with a C-terminal polyhistidine sequence as previously described [30], and a metal affinity chromatography purification step was substituted for anion exchange chromatography. The F433A mutant was generated by PCR mutagenesis with this expression construct and sequenced to verify the change.

### PSA crystallization and structure determination

PSA was crystallized by hanging drop vapor diffusion with initial conditions defined using commercially available solution screens (Hampton Research; Molecular Dynamics Ltd.). High quality crystals were obtained reproducibly at 16°C using 5-8 mg/ml PSA mixed 1:1 with 15% PEG 4K, 0.1 M Tris (pH 8.5), 0.5 M sodium chloride, and 1.5% v/v dioxane. PSA crystals generally grew to full size in two weeks.

Crystals were prepared for X-ray data collection by serial transfer through solutions containing the crystallization components plus glycerol at concentrations increasing from 5 to 30% in 5% steps. Crystals were soaked in each solution for approximately 10 minutes. The crystals were then mounted in nylon loops and flash cooled by plunging into liquid nitrogen [36]. Data were collected on beamline 22ID at the Advanced Photon Source, Argonne National Laboratory and processed with HKL2000 [37]. The structure was determined by molecular replacement using tricorn interacting protease factor F3 (PDB ID 1Z1W) [38] as a search model. The search model is 47% identical or conserved over the PSA sequence used for crystallization. Automated model building was carried out with Phenix [39, 40], followed by iterative rounds of manual building in COOT [41] and refinement in Phenix. Molecular figures were produced using PYMOL (http://www.pymol.org). The final structure contains one dioxane and one Tris ligand in addition to ordered solvent.

### Polyglutamine peptide purification

A commercially prepared (Peptidogenic Research and Co., Livermore, CA) polyglutamine peptide (PQ) with the sequence Lys_2_Gln_15_Lys_2_ was purified with slight modifications to a published protocol [42]. 5 ml of trifluoroacetic acid and 5 ml of hexafluoro-2-propanol (TCI America) was added to a glass vial containing 4 mg of the PQ peptide. The mixture was vortexed intermittently for 2 minutes and left overnight at room temperature. The solvent was evaporated over a period of one hour using a gentle stream of argon gas, and then placed on a lyophilizer for half an hour to remove any residual solvent. Then 1 ml water (made to pH 3.0 with TFA) was added to the sample. This sample was spun at 50,000 x *g* for 3.5 hours at 4°C to separate any remaining aggregated material. The top two thirds of the supernatant was recovered and flash frozen in liquid nitrogen for later use.

### PSA-PQ complex

PSA protein crystals were grown as described above. The crystals were transferred into a solution containing 1mM EDTA, 0.1 M Tris (pH 8.5), and 0.5 M sodium chloride and left to soak for 1 hour. These crystals were then transferred for 45 min into a solution containing the crystallization conditions plus 1mM EDTA and 200 µM PQ peptide. The crystals were flash cooled by transferring them briefly to a cryosolution containing crystallization conditions plus 1mM EDTA, 200 µM PQ peptide and 20% glycerol and then plunging loop mounted crystals into liquid nitrogen. Data were collected and processed as described above, resulting in a complete 3.65 Å data set. Difference maps were generated using the Phenix package [39] with the unliganded PSA structure, and poly-alanine was modeled into the observed difference density with COOT [41] using manual building and real space refinement for the peptide only. Subsequently, the polyalanine peptide was converted to polyglutamine, and the PSA-PQ complex was refined in Phenix using positional restraints to the unliganded PSA structure because of the limited data resolution. Since there was generally no convincing electron density for the side chains beyond the beta carbons, the peptide was converted to polyalanine for deposition after an additional round of refinement.

### Kinetic analysis

The kinetic properties of PSA were determined using the fluorogenic substrate alanine 4-methoxy-β-naphthylamide (Ala-4MβNA) in 20mM HEPES pH7.0 and 2 mM BME at 37°C [10]. Release of free naphthylamide was monitored on a fluorescent plate reader (SpectraMax Gemini XS) at an excitation wavelength of 335 nm and an emission wavelength of 410 nm. In particular, estimates of the K_i_ values for the PQ peptide and dynorphin A(1-17) were determined based on inhibition of Ala-4MβNA cleavage [10] at increasing concentrations of either peptide. Reactions were carried out with and 20 µM (PSA^wt^) or 100 µM (PSA^F433A^) Ala-4MβNA in a total volume of 200 µl, with reaction rates measured in triplicate. Single reciprocal plots of the inhibition data were fit by linear regression using the Prism software package (GraphPad Prism), with the X intercept representing K_i_(1+[S]/K_m_) where [S] is the concentration and K_m_ is the Michaelis constant of Ala-4MβNA. K_m_ values for Ala-4MβNA were 29.2 µM for PSA^wt^ and 160 µM for PSA_F433A._

## Results

### PSA architecture

Data and model statistics for crystal structures of PSA and PSA with bound peptide are provided in **Table 1**. PSA adopts a lopsided V-shape overall conformation, creating a central groove that is about 20 Ǻ long and 15 Ǻ wide (**Fig 1**; secondary structure versus sequence and residue number shown in **Fig 2**). The longer arm of the V consists of residues from the N terminus through residue 594, while the shorter, C-terminal arm comprises residues 595-914. The overall architecture is similar to other M1 family peptidases with known structures, including tricorn interacting factor F3 [38], aminopeptidase N from *Escherichia coli* (ePepN) [43, 44], aminopeptidase from *Plasmodium falciparum* (Pfa-M1) [45], and the smaller leukotriene A4 hydrolase (LTA4H) [46]. More recently, the human endoplasmic reticulum aminopeptidases (ERAP1 and ERAP2) have also been shown to share the same overall fold [47–49]. The N-terminal arm of PSA can be further divided into 3 distinct regions: N-terminal domain (domain I, residues 51-253; residues 1-50 not ordered in the crystal structure), catalytic domain (domain II, residues 254-503), and linker domain (domain III, residues 504-594). This arrangement is primarily defined by the presence of the central metallopeptidase domain, which has strong structural similarity to the bacterial metalloprotease thermolysin [50, 51].

**Figure 1.**
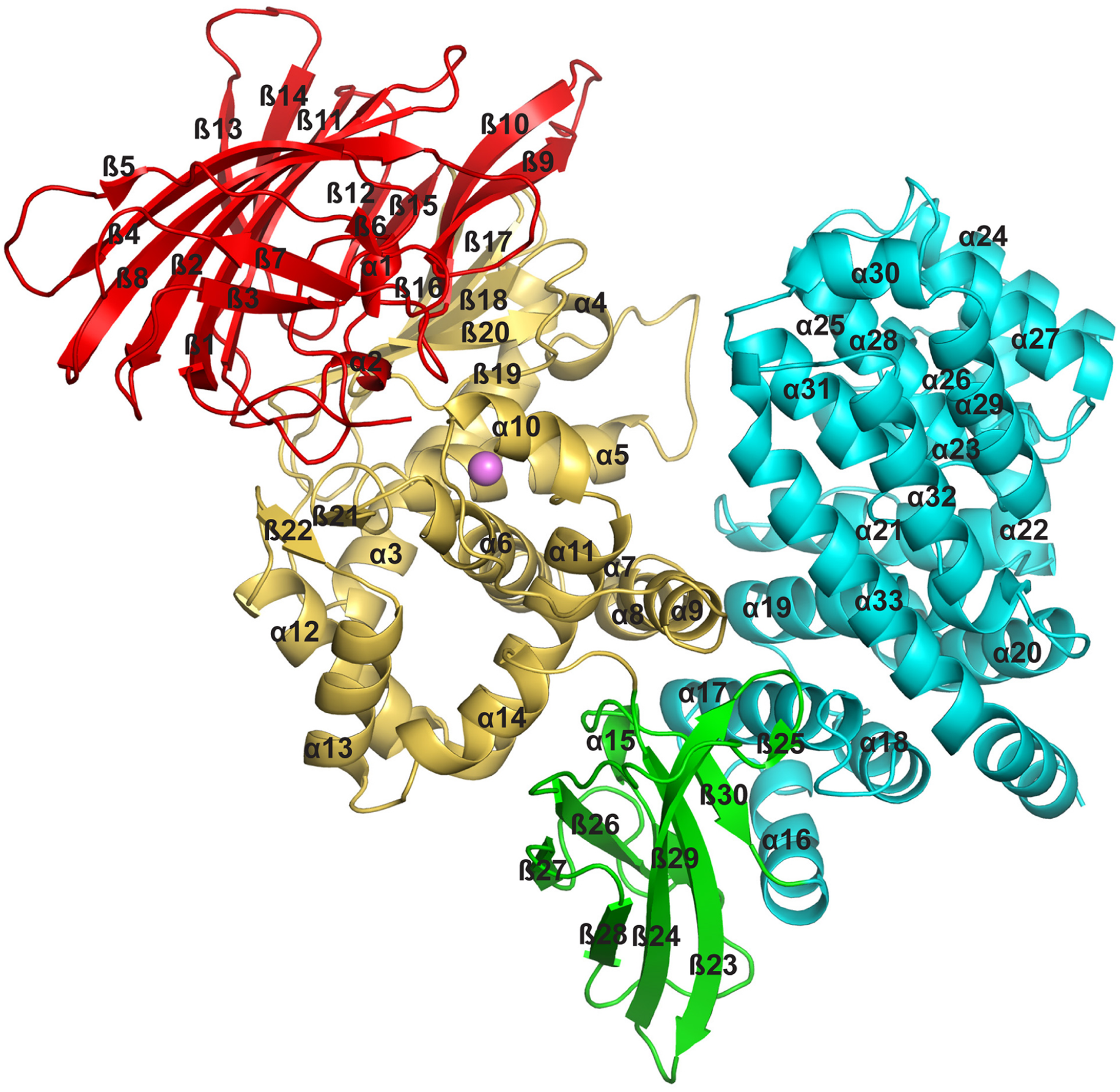
Overview of the PSA structure. The backbone of PSA shown in a ribbons representation with domains in different colors: N-terminal domain I (red), catalytic domain II (gold), linker domain III (green), C-terminal domain IV (cyan). Secondary structure elements are labeled, and the active site zinc ion is shown as a pink sphere.

**Figure 2.**
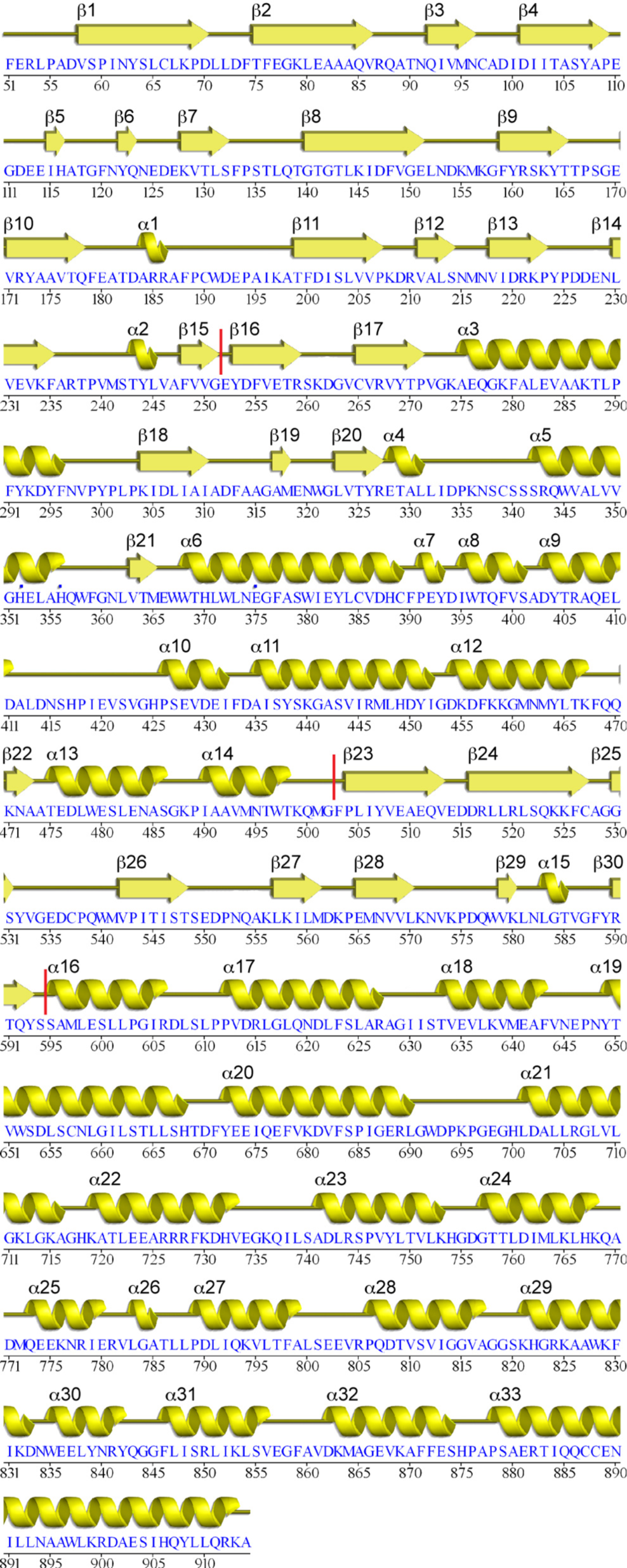
PSA sequence and secondary structure. Sequence and corresponding secondary structure of human PSA are illustrated. Vertical red lines indicate domain boundaries.

**Table 1.** Crystallographic statistics.

|  | PSA | PSA-PQ peptide |
| --- | --- | --- |
| **Data collection** |  |  |
| Space group | P2<sub>1</sub>2<sub>1</sub>2 | P2<sub>1</sub>2<sub>1</sub>2 |
| Cell dimensions |  |  |
| *a*, *b*, *c* (Å) | 70.7, 257.7, 60.4 | 70.7, 256.6, 60.3 |
| $\alpha$, $\beta$, $\gamma$ (°) | 90.0, 90.0, 90.0 | 90.0, 90.0, 90.0 |
| Resolution (Å) | 50-2.3 (2.38-2.30)<sup>a</sup> | 50-3.65 (3.84-3.65) |
| *R*<sub>merge</sub> | 0.065 (0.587) | 0.133 (0.567) |
| *R*<sub>pim</sub> | 0.033 (0.306) | 0.079 (0.346) |
| CC<sub>1/2</sub> in highest shell<br>(number of pairs) | 0.75 (2279) | 0.697 (938) |
| $I/\sigma I$ | 20.7 (2.4) | 8.2 (2.0) |
| Completeness (%) | 95.9 (95.4) | 99.2 (99.7) |
| Redundancy | 4.7 (4.6) | 3.6 (3.6) |
| **Refinement** |  |  |
| Resolution (Å) | 45.2-2.3 (2.38-2.30) | 47.5-3.65 (3.76-3.65) |
| Total reflections | 47,918 | 12,765 |
| Reflections in refinement | 47,914 | 12,726 |
| $R_{\text{work}}$ | 0.223 (0.321) | 0.257 (0.361) |
| $R_{\text{free}}$ | 0.241 (0.335) | 0.295 (0.418) |
| Number non-hydrogen atoms | 7055 |  |
| Protein (one in a.u.) | 6856 | 6856 |
| Zn<sup>2+</sup>, | 1 | 1 |
| Dioxane, Tris | 14 | 0 |
| Water | 184 | 0 |
| Protein residues | 864 | 864 |
| Peptide residues | 0 | 6 |
| Average B-factors (Å<sup>2</sup>) |  |  |
| Protein | 68 | 131 |
| Domain I<sup>b</sup> | 90 | 143 |
| Domain II | 64 | 125 |
| Domain III | 51 | 109 |
| Domain IV | 61 | 135 |
| 1<sup>st</sup> HEAT repeat | 50 | 117 |
| 2<sup>nd</sup> HEAT repeat | 71 | 149 |
| Peptide |  | 204 |
| Zn<sup>2+</sup>, | 37 | 140 |
| Dioxane, Tris | 70 |  |
| Water | 53 |  |
| Number of TLS groups | 3 |  |
| R.m.s deviations |  |  |
| Bond lengths (Å) | 0.002 | 0.002 |
| Bond angles (°) | 0.42 | 0.44 |
| Ramachandran plot (%) |  |  |
| Favored | 95.6 | 95.2 |
| Allowed | 4.2 | 4.5 |
| Outliers | 0.2 | 0.3 |

<sup>a</sup> Values for the last resolution shell are shown in parentheses.
<sup>b</sup> Domain I (residues 51-253), domain II (residues 254-503), domain III (residues 504-594), domain IV (residues 595-914). First HEAT repeat (residues 595-739), second HEAT repeat (residues 740-914).

Domain I of PSA is dominated by a large beta sheet that forms the end of the longer arm of the molecule (**Fig 3A**). The eight strands in this sheet are arranged in a mixed orientation and adopt a saddle shaped structure. Three smaller sheets of three, two, and two strands, tuck under the ends of the saddle, where they form part of the interface with the catalytic domain. The eight-stranded sheet consists of beta strands 1, 2, 4, 5, 8, 11, 13 and 14. The three-stranded sheet (strands 3,6 and 7) has antiparallel strands with one solvent exposed face, and the two stranded sheets (strands 9 and 10 and strands 12 and 15) are nearly parallel and in the same plane at the other end of the saddle. These two smaller sheets form a beta sandwich element with a sheet from the catalytic domain, making extensive contacts at the interface. The N-terminal domain of PSA appears to be unique to the M1 aminopeptidases, with no strong similarities detected to structures outside the family.

**Figure 3.**
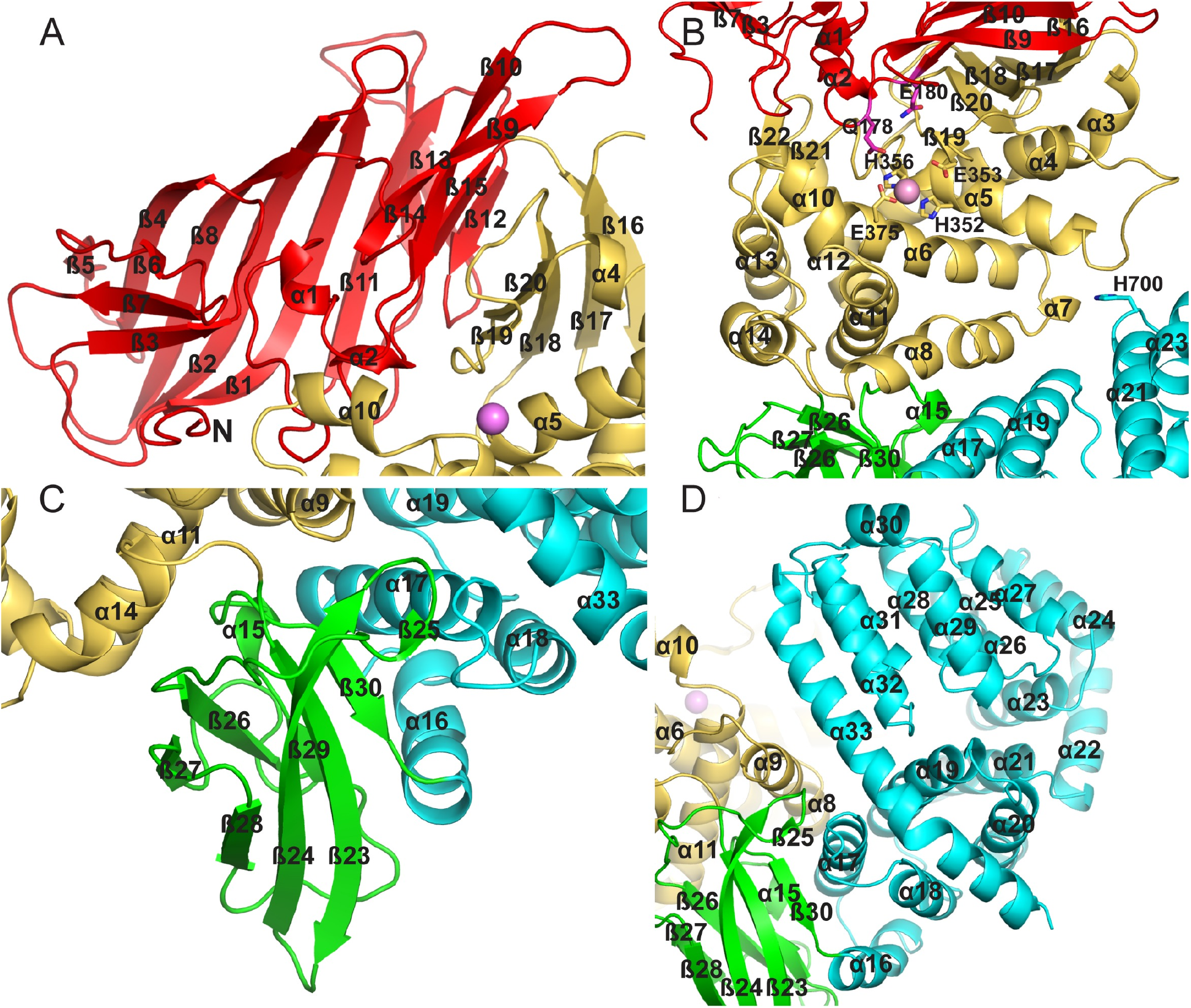
PSA domain structure and domain interactions. Overview of the individual domain structures and interfaces for the (A) N-terminal (Domain I), (B) catalytic (Domain II), (C) linker (Domain III), and (D) C-terminal (Domain IV) domains. Secondary structural elements are labeled, as are residues mentioned in the text.

The catalytic domain (domain II) is composed of a mixed β-sheet and a large helical cluster consisting of helices 3-14 with a small two-stranded parallel sheet in addition (**Fig 3B**). The mixed sheet consists of strands 16-20, and it forms a large part of the interface with the N-terminal region. All zinc metallopeptidases with known structures that function as neuropeptidases have a conserved thermolysin-like [52–55] active-site fold [56], and the active site domain of PSA superimposes on thermolysin (PDB ID 1L3F) with an r.m.s.d. of 5.2 Å over 216 Cα atoms out of 251 residues. The active site itself contains the conserved motif HEXXH, which is present in α5. As in other metallopeptidases, the two histidine residues (His352 and His356) of the motif coordinate the zinc ion, and the glutamate residue (Glu353) hydrogen bonds to a water molecule that is also coordinated to the metal (**Fig 4**). In addition, another glutamate residue (Glu375) present in α6 acts as a fourth zinc-coordinating group. This coordination of the zinc ion by two histidine residues and a downstream glutamate is characteristic of clan MA metallopeptidases. In PSA and other M1 aminopeptidases, the downstream glutamate is part of a conserved sequence motif, NEXFA [38, 43, 50, 51].

**Figure 4.**
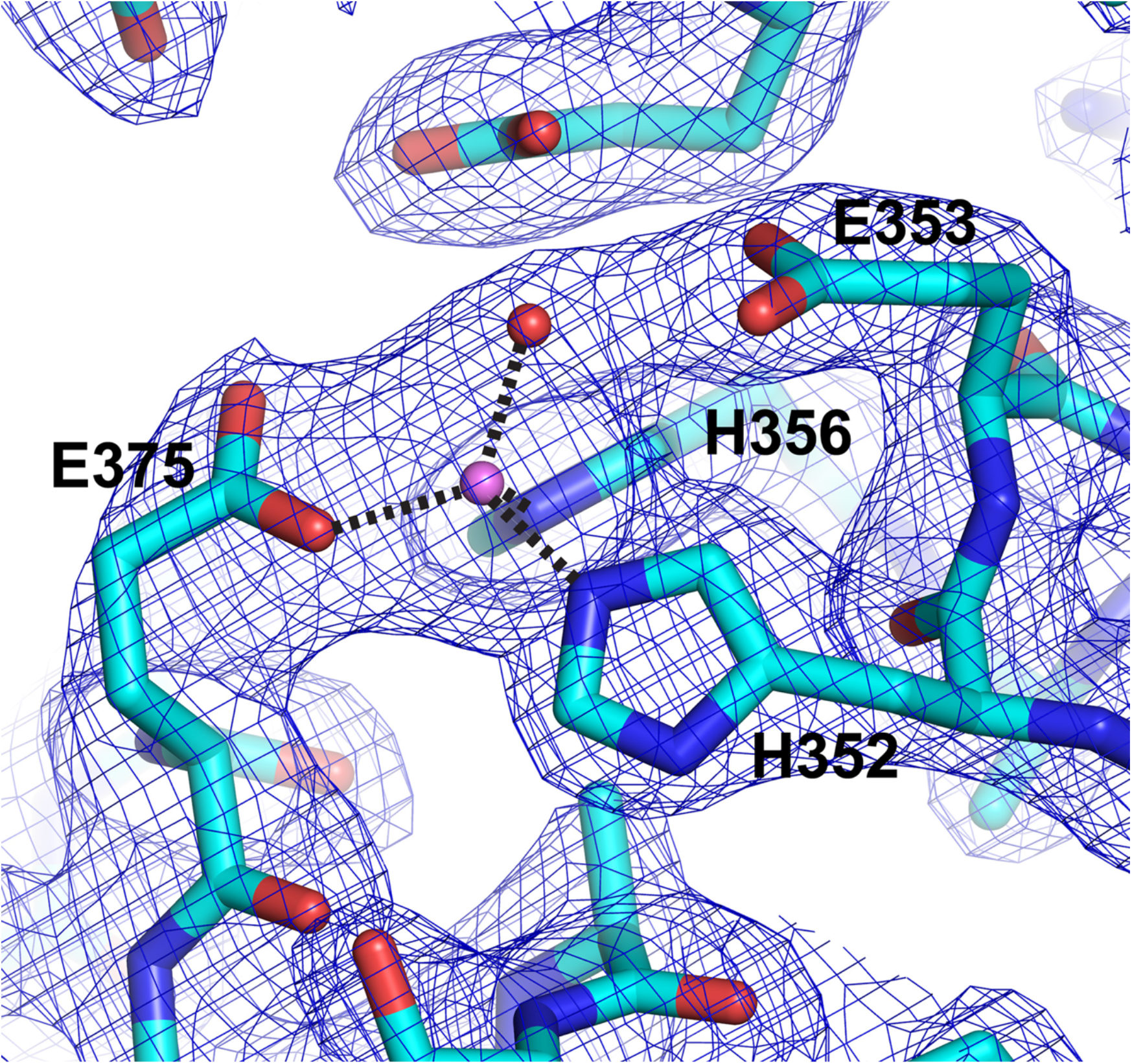
PSA active site. The active site of PSA is shown in a stick representation with the catalytic zinc ion and water molecule indicated by the pink and red spheres, respectively. Electron density (2Fo-Fc, 1.2 σ contour) is shown in blue. Zinc coordinating residues and the catalytic glutamate are labeled. Coordination interactions with the zinc ion are shown as dashed lines. In the crystal structure, the coordination distances to the zinc ion are: E375, 1.96 Å; H352, 2.12 Å; H356, 2.11 Å; and the coordinating solvent, 2.03 Å.

Residues from helices 6, 8, 9,11 and 19 largely make up the floor of the active site. The active site walls are formed by the edge of the five-stranded sheet (strand 16), helices 5, 10, 21, 25, 28, 31 and 33 as well as loops made up of residues 151-159 and 327-340. While the active site is principally comprised of residues from the catalytic domain, the N-terminal domain contributes residues, including Gln178 and Glu180, and helices from the C-terminal domain (domain IV) form one side of the active site pocket. In some conformations of factor F3, an arginine residue (Arg721) from domain IV contacts an extended loop (residues 324-250) from the active site domain, particularly interacting with the side chain of Phe346 [38]. In PSA, the equivalent of Arg721 is Phe846, but the segment containing this residue (around the N terminus of α31) is shifted away from the catalytic domain by about 6 Å relative to factor F3, and no contacts are made. There are interactions, however, between the N terminus of α5, which contains the HEXXH motif, and the turn between helices 20 and 21 (particularly His700) in the C-terminal domain. Interactions also occur between helices 8 and 9 of domain II and α17 from domain IV. Although neither of these interactions is extensive, they likely contribute to maintaining the open conformation of the enzyme.

The linker domain (domain III) serves to make the connection between the N- and C-terminal portions of the enzyme (**Fig 3C**). This domain follows the metallopeptidase active site region and consists of two sheets containing strands 23, 24, 25, 28, and 30, and 26, 27 and 29 that are packed against each other to form an immunoglobulin-like β-sandwich fold. The smaller three-stranded sheet primarily makes contacts with the domain II (particularly α11 and α14), while the edges of both sheets interact with domain IV (α16 and α17). The interface with domain II buries 2010 Å^2^ of solvent exposed surface, while the contact with domain IV is about the same size, burying 1990 Å^2^ of exposed surface. Both domain-domain interfaces are largely hydrophobic and aromatic in nature, and it seems likely that they are rigid and stable.

The C-terminal domain IV forms the short arm of the V-shaped PSA molecule (**Fig 3D**). It consists of 18 helices arranged into two superhelical HEAT repeat segments [57]. The first six helices (α16-21) form one superhelical segment. Two subsequent helices (α22 and α23) serve to turn the path of the superhelix roughly 120°, and the remaining ten helices (α24-33) form the second superhelical segment. The C-terminal helix (α33) of this second segment is elongated, and it interacts with the first superhelical segment to form a closed loop. Domain IV is known to be required for proper folding of the remainder of the molecule when expressed in *E. coli* [58]. The conformation of the entire domain is unique to the M1 peptidases, but as expected, a number of other proteins with HEAT repeats show structural similarity to the individual repeats of PSA, particularly the longer C-terminal repeat. The C-terminal domain in endoplasmic reticulum aminopeptidase 1 (ERAP1) has been shown to interact with the C terminus of a bound 15-residue peptide analog [59]. The interacting residues are not conserved in PSA, but other residues in the corresponding region, particularly Lys712 and Lys715, might mediate a similar interaction. More generally, domain IV of ERAP1 has been shown to mediate binding of the C termini of peptide-like inhibitors, as well as an allosteric effector, at a distributed set of sites, which can serve to modulate a large-scale conformational change in the enzyme [60, 61]. PSA domain IV could play a similar role.

HEAT repeat superhelices often mediate protein-protein interactions [57]. PSA has been reported to co-localize with tubulin [11, 62], and it is interesting to note that one of two putative microtubule associated protein (MAP) sequences present in PSA [11] is located within the first superhelical segment of the domain IV (residues 682-703; **Fig 5**). In Tau and a number of other proteins that interact with microtubules, MAP sequences help to mediate the binding interaction [63–67]. The other putative MAP motif in PSA is located in the catalytic domain (residues 266-289), where it comprises the C-terminal portion of β17, the following loop segment, and the N-terminal portion of α3. In contrast, the MAP motif in domain IV forms portions of two helices: the C-terminal segment of α20 and the first turn of α21, as well as the nine-residue intervening loop, which contains a sequence similar to the most conserved Pro-Gly-Gly-Gly sequence of the MAP motif. In MAP sequence containing proteins, the motif appears to be unstructured when not interacting with tubulin, but a portion of the sequence may form an additional strand or two of an α-tubulin sheet when bound to microtubules as seen in the crystal structure of a complex between tubulin and the MAP-related stathmin-like domain sequence [66, 68]. While the PSA MAP sequence in domain II has only a relatively short loop segment that is not positioned well to interact with a large structure like a microtubule, the longer loop in the domain IV MAP sequence points into solvent in an orientation that would likely allow it to mediate an interaction with microtubules. In that regard, some proteins that promote tubulin polymerization interact via TOG domains, which are formed from HEAT repeats [69–72], and it is possible that other loop segments in the domain IV HEAT repeats contribute to an interaction with microtubules. The two MAP sequences in PSA also show similarity to a characteristic sequence motif in some proteasome subunits [11]. For example, the proteasome sequence occurs near the N-termini of the alpha subunits in the yeast 20S proteasome structure, forming an open coil segment and a short helix that make up part of the gate assembly of the proteasome [73]. While this structure of the motif is like the one adopted by the domain II MAP sequence, no functional significance of this similarity is suggested by existing knowledge of PSA activity.

**Figure 5.**
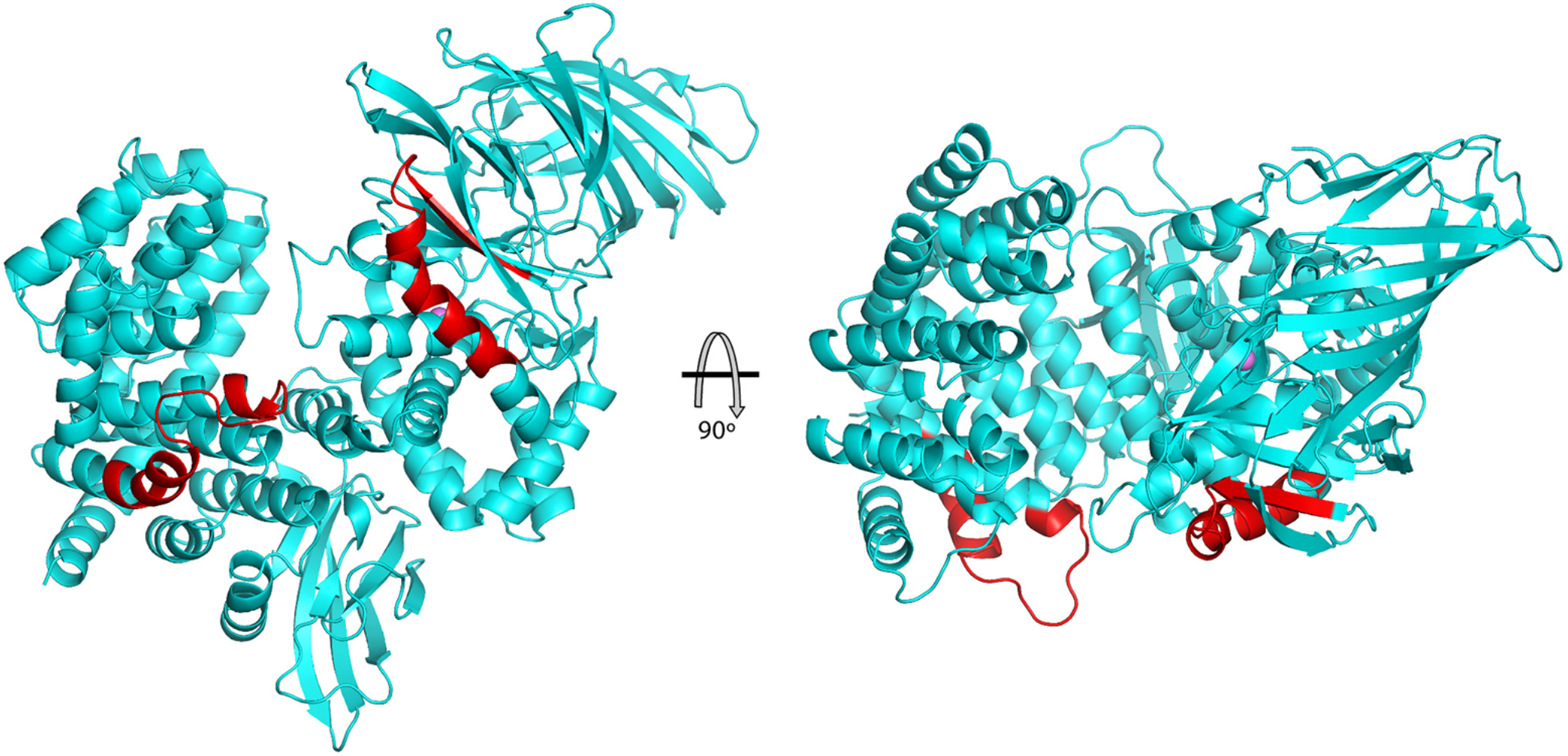
Microtubule associated protein sequences in PSA. The location of the microtubule associated protein sequences are shown in red on a ribbons trace of the PSA structure. One sequence is in the catalytic domain (domain II) and the other in the C-terminal domain (domain IV). The two views are related by a 90° rotation.

### Comparison of M1 aminopeptidases

The crystal structures of other aminopeptidases in the M1 family include: aminopeptidase A (APA) [74], tricorn interacting factor from *Thermoplasma acidophilum* (factor F3) [38], aminopeptidase N (APN) [75–77] and leukotriene A4 hydrolase (LTA4H) [46], *P. falciparum* aminopeptidase (PfA-M1) [45], aminopeptidase N from *E. coli* (ePepN) [43, 44], endoplasmic reticulum aminopeptidase 1 (ERAP1) [47, 48], endoplasmic reticulum aminopeptidase 2 (ERAP2) [49], aminopeptidase N from *Anopheles gambiae* (AnAPN1) [78], insulin regulated aminopeptidase (IRAP) [79, 80], cold-active aminopeptidase from *Colwellia psychrerythraea* (ColAP) [81], aminopeptidase A from *Legionella pneumophila* (LePepA) [82] and aminopeptidase N from Deinococcus radiodurans (M1dr) [83, 84]. Alignment of the human PSA sequence with these other M1 members indicates they are not closely related. Sequence identities range from 15-32% (similarity 24-49%), with most of the human paralogs (APA, APN, ERAP1, ERAP2, and IRAP), as well as factor F3 and AnAPN1, being at the high end of the range. Although the lopsided V-shaped overall architecture of PSA is maintained in the other aminopeptidases, most of the enzymes crystallize in a closed conformation where a portion of domain IV shifts to interact with the catalytic domain II, eliminating the gap between the two arms of the V (**Fig 6**). This conformational change primarily involves a rigid rotation of the domain IV second HEAT repeat (and the transition helix α22).

**Figure 6.**
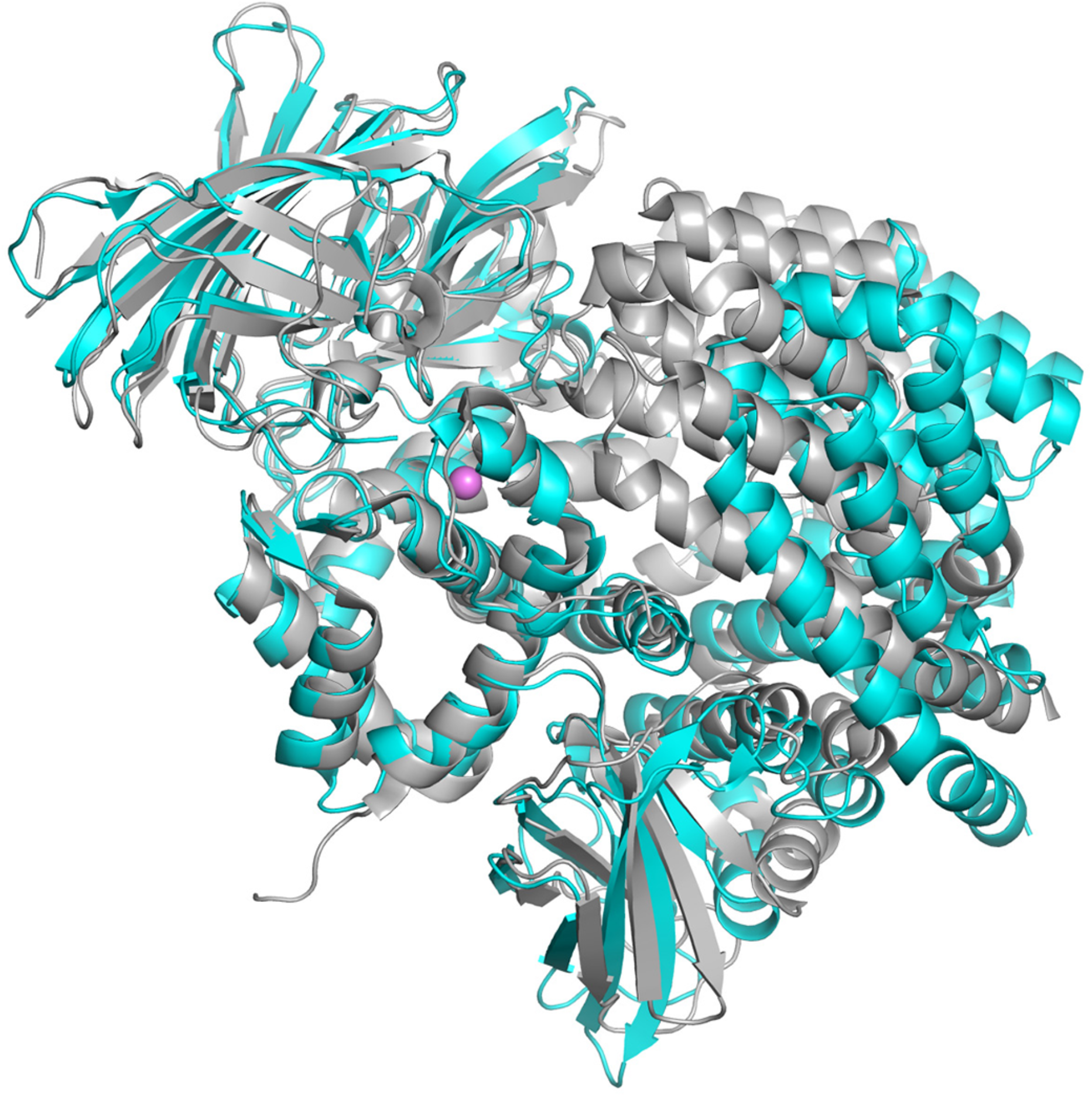
Open and closed conformations. A ribbons view of PSA (cyan) is shown superimposed on ERAP2 [49] (gray; PDB ID: 3SE6). The more open conformation of PSA, with domains II and IV separated by a wide channel, results in a larger and more accessible active site region.

Several crystal forms of ERAP1 [47, 48, 61], two forms of APN [75, 76], and factor F3 [38] also adopt the fully open conformation seen in PSA. (Unliganded IRAP is not open to the same extent as PSA, but domain IV is not in contact with domain II. Binding of an inhibitor causes it to adopt a fully closed conformation [85].) The enzymes in the closed conformation generally have been crystallized with bound inhibitors, peptides, or individual amino acids at the active site. However, it appears that crystallization may trap lower probability conformers, for example unliganded Pfa-M1 in the closed form [45] or ERAP1 bound to peptide-like inhibitors in the open form [48, 61]. In the case of ERAP1, X-ray scattering and other studies convincingly demonstrate that substrate mimics shift the conformational equilibrium toward the closed form in solution [60, 61], and this seems likely the case for at least most other M1 members. The AlphaFold Database prediction for human PSA adopts the closed conformation [86, 87], which may be influenced by the abundance of closed conformation templates. Running AlphaFold Colab [86] without templates, however, also generates a prediction in the closed conformation, indicating coevolutionary restraints at the domain II-IV interface. The closed PSA model has an internal chamber with only narrow solvent openings. CASTp [88] reports a volume of 3510 Å^3^ for this chamber, which would be sufficient to accommodate the largest reported PSA substrate, dynorphin A(1-17) with an excluded volume of 2023 Å^3^. Except for the shift of domain IV, the individual domains of all the M1 peptidases maintain the general structure seen in PSA with RMSD values on Cα superposition varying between 1.2 to 4.3 Å. Exceptions are LTAH4 and ColAP, which lack the linker domain entirely and have a single HEAT repeat in domain IV [46, 81], as well as M1dr, which has only the N-terminal and catalytic domains [83, 84]. The backbone conformations of the AlphaFold PSA model individual domains agree well with those of the PSA crystal structure, giving Cα superposition RMSD values of: domain 1, 0.42 Å; domain 2, 0.75 Å; domain 3, 0.35 Å, domain 4, 0.66 Å.

Notably, average atomic thermal factors are higher in PSA domain I and the second HEAT repeat of domain IV (see **Table 1**). Since the largest difference between the open PSA conformation and the closed conformation of other M1 aminopeptidase structures is a shift of the domain IV second HEAT repeat, the increased thermal factors in that PSA domain may reflect a true increase of dynamics or flexibility of this region relative to the remainder of the molecule. Some other aminopeptidase structures crystallized in open conformations, F3 (PDB ID 1Z1W) and ERAP1 (PDB ID 3MDJ, 3QNF), and APN (PDB ID 4FKE, 4F5C), also show higher thermal factors for the second HEAT repeat of domain IV.

### Other residues involved in catalysis

Stabilization of the oxyanion generated in the transition state by zinc metallopeptidases generally involves not only the positively charged zinc ion but also one or two hydrogen bond donating side chains [50]. In thermolysin, His231 and Tyr157 likely donate hydrogen bonds in this manner. Tyr438 in PSA, which is in α11, occupies the position equivalent to His231 and could participate in transition state stabilization (**Fig 6A**). Mutating this residue to phenylalanine reduces k_cat_ by1000 fold, indicating its importance in catalysis [35]. Tyr157 in thermolysin is in the loop between the active site helices. The most structurally equivalent residue in PSA is Trp367, which is conserved in a number of other M1 aminopeptidases. This residue is far (over 16 Å) from the active site zinc ion, however, and is unlikely to participate in catalysis. Tyr378 in LTA4H has been proposed to act as a second stabilizing residue [46]. This tyrosine is conserved in APN [43] but it is replaced by phenylalanine in both PSA and Factor F3 and therefore could not function to stabilize the oxyanion in these enzymes. Another tyrosine, residue 244, is in the vicinity of the active site in PSA where it could possibly participate in catalysis (see **Fig 7A**). It is located in the turn connecting strands 14 and 15 of the N-terminal domain, however the distance between its hydroxyl group and expected position of the carbonyl oxygen is over 8 Å, indicating that a conformational change would be needed for it to participate in transition state stabilization.

**Figure 7.**
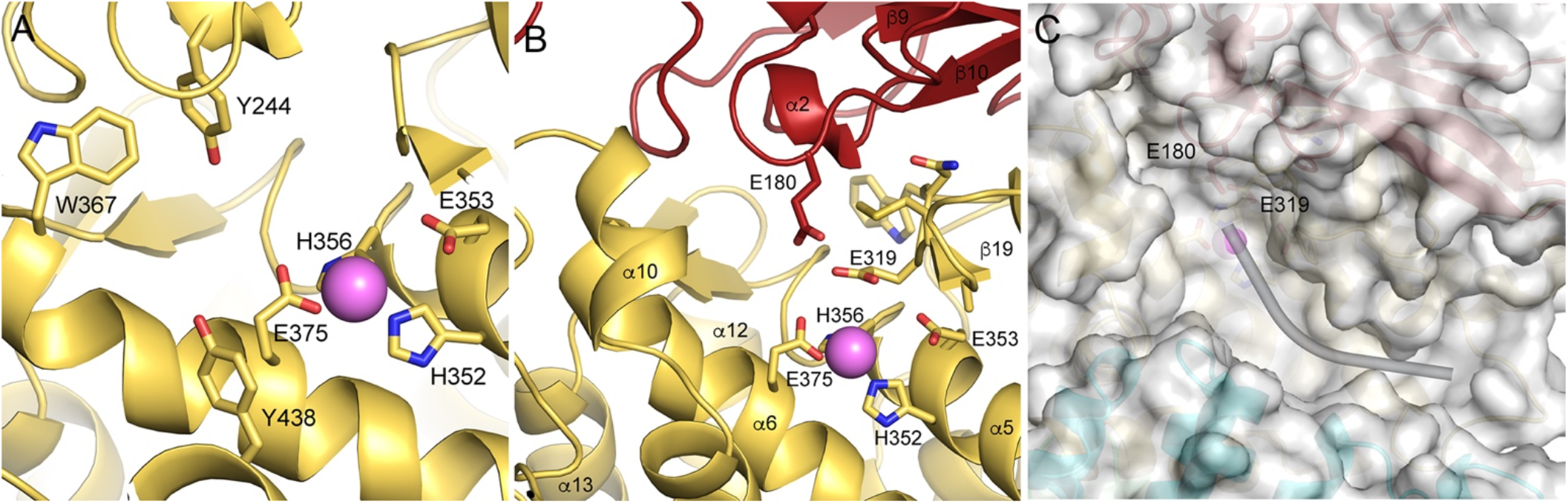
Active site features. (A) Active site region of PSA showing side chains of zinc coordinating residues and other residues that may participate in catalysis as discussed in the text. The active site zinc ion is shown as a pink sphere. (B) View of the active site with elements that close off one end to promote aminopeptidase activity. Side chains for the conserved Glu-Ala-Met-Glu-Asn-Trp (GAMENW) sequence are shown in a stick representation. The Glu180 side chain from the N-terminal domain (red) is also shown. Glutamates 180 and 319 likely interact with the N terminus of bound peptide substrates. (C) Molecular surface (semi-transparent) and ribbons view of the active site region. Glu319 of the GAMENW sequence is indicate, as is nearby Glu180. A likely path for bound peptide substrate is indicated by the curved cylinder. The active site zinc ion is in pink.

### Aminopeptidase activity

Thermolysin and many other zinc metallopeptidases act as endopeptidases, cleaving peptide sequences internally. Their active sites allow bound substrate peptides to extend in either direction from the catalytic machinery. In PSA, however, the active site is closed off at one end by elements of the N-terminal domain (**Fig 7B,C**), restricting the extent of the peptide N-terminal to the cleavage site. Specifically, strands and turns from the second β sheet of that domain, as well as residues from the turns between helices 5 and 6 and helices 12 and 13 pack to form a structural wall that limits substrate binding to one amino acid N-terminal to the scissile bond. Thus, the active site channel of PSA resembles a blind canyon with the catalytic machinery located near its closed end. The aminopeptidase specific GAMENW sequence [43, 44, 89, 90] encompasses one edge strand (β19; residues 316-321) of the five-stranded sheet in the catalytic domain and the following open coil segment. Glu319 from this sequence is well positioned to interact with the N-terminal amino group of bound peptides, as has been proposed [43, 44]. Together with the nearby Glu180, it creates a pocket with strongly negative electrostatic potential that likely binds the N-terminus of substrate peptides, helping to position them appropriately for catalytic removal of the first residue. Near the active site, bound substrate peptide likely interacts with main chain groups of residues in strand 19, the edge of the catalytic domain central sheet, as expected for the binding of substrates to zinc metallopeptidases [56]. In particular, the carbonyl group of Ala317 likely accepts a hydrogen bond from the main chain amine of the P1 substrate residue. In addition, the main chain amine of Ala317 is in position to interact with the carbonyl group of the P1’ substrate residue. The side chains of Val349, Ser379, and Glu382 are also in position to interact with the P1’ residue depending on the path of the substrate as it exits the immediate active site region.

The active site of PSA lies at the end of a long groove in the enzyme, presenting a large surface that likely provides the basis for interaction with the extended portion of peptide substrates C terminal to the scissile bond (**Fig 8**). Interestingly, this potential substrate-binding surface in PSA is enriched in aromatic and hydrophobic residues, with some 36 solvent-exposed hydrophobic/aromatic residue side chains around the active site. The floor and sides of the active site channel are lined with these hydrophobic/aromatic residues, and, in particular, residues from helices α5 and α6 make much of the floor of the channel where substrates likely interact.

**Figure 8.**
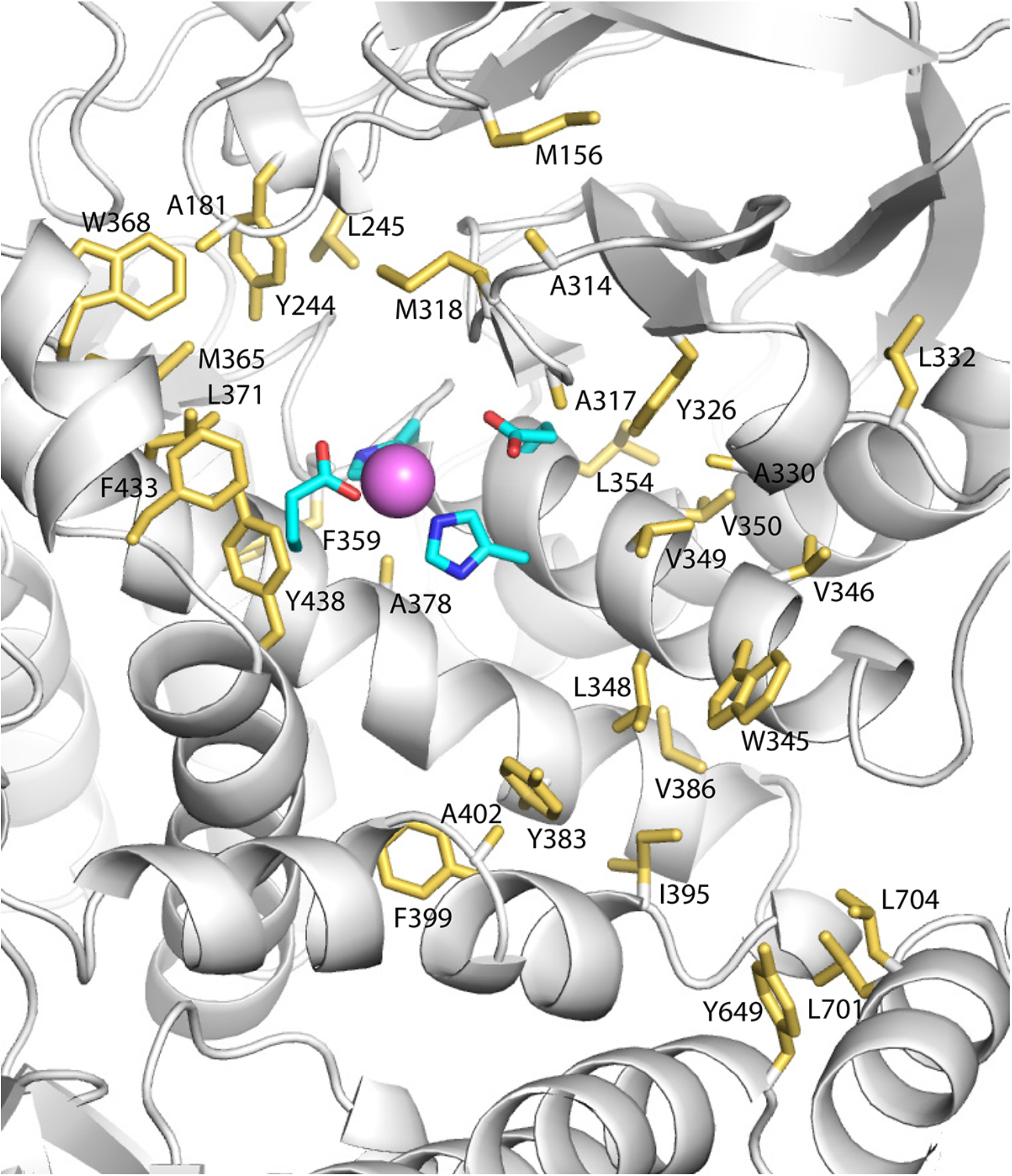
Extended peptide-binding surface in PSA. Side chains of hydrophobic and aromatic residues that form parts of the molecular surface surrounding the active site of PSA are shown in a gold stick representation. The active site zinc ion (pink sphere) and side chains of active site residues are also shown.

### Hydrolysis of polyglutamine peptides

Glutamine-rich sequences are found in many cellular proteins, including those associated with neurodegenerative disorders. The Huntingtin protein, for example, contains polyglutamine sequences [91] prone to expansion as a result of errors during DNA replication [92]. Expanded polyglutamine sequences tend to aggregate [93] and form inclusions that are the pathological hallmarks of neurodegenerative diseases like Huntington’s disease and spinocerebellar ataxia [92, 94]. Importantly, polyglutamine tracts are not degraded efficiently by the proteasome [95]. PSA is the only proteolytic activity in HeLa cells identified as being responsible for processing polyglutamine sequences, and the enzyme was able to degrade long polyglutamine peptides (20-30 glutamines) as efficiently as short polyglutamine containing peptides [31]. Moreover, PSA knockdown or inhibition increased polyglutamine accumulation and PSA overexpression had the opposite effect in other cell types [32]. To help identify specific residues in PSA that may have a role in polyglutamine turnover, we determined the crystal structure of PSA in complex with a 19-mer polyglutamine peptide (PQ) having the sequence Lys_2_ - Gln_15_ -Lys_2_.

Difference electron density in the active site region (**Fig 9A**) defined the binding site of the PQ peptide, and a polyalanine peptide was initially modeled into the low resolution density. The bound PQ is in position to interact with the glutamate residue (Glu319) of the GAMENW aminopeptidase recognition sequence in β19 as well as Glu319 from domain I. The peptide initially extends along strand 19, donating a hydrogen bond from the P1 residue to the carbonyl group of Ala317. The path of the peptide then turns, however, allowing it to interact with residues in helix 10 and the following loop, which includes Phe433. Fitting the backbone density in this manner results in a cis peptide bond between the P1 and P1’ residues, but this unusual configuration may be a result of uncertainty in the build due to the low resolution of the density. Subsequent restrained refinement of the peptide converted to polyglutamine showed little side chain density with the exception of the P4’ residue (**Fig 9B**). Nevertheless, the glutamine side chains can adopt favorable conformations with no major clashes, and their positions suggest potential interactions with the protein. The P1 sidechain may interact with Glu375 and Tyr438, and Gln178 is positioned to contact the side chain of the P1’ residue. The side chain of the P3’ residue may interact with Phe433 and the P4’ side chain with nearby Asp430. The electron density after P5’ becomes weak, preventing any further tracing of the substrate backbone path. In all, 6 alanine residues were included in the final peptide model. Interestingly, the electron density indicates that the zinc ion cofactor was present with high occupancy in the crystal despite the EDTA soak intended to remove it. Thus, the enzyme would have retained at least partial activity during peptide soaking, and it is likely that the bound fragment represents an average of partially degraded peptides of different lengths. The electron density is consistent with the carbonyl oxygen of the first substrate peptide bond coordinating the zinc ion in an orientation similar to that expected during hydrolysis.

**Figure 9.**
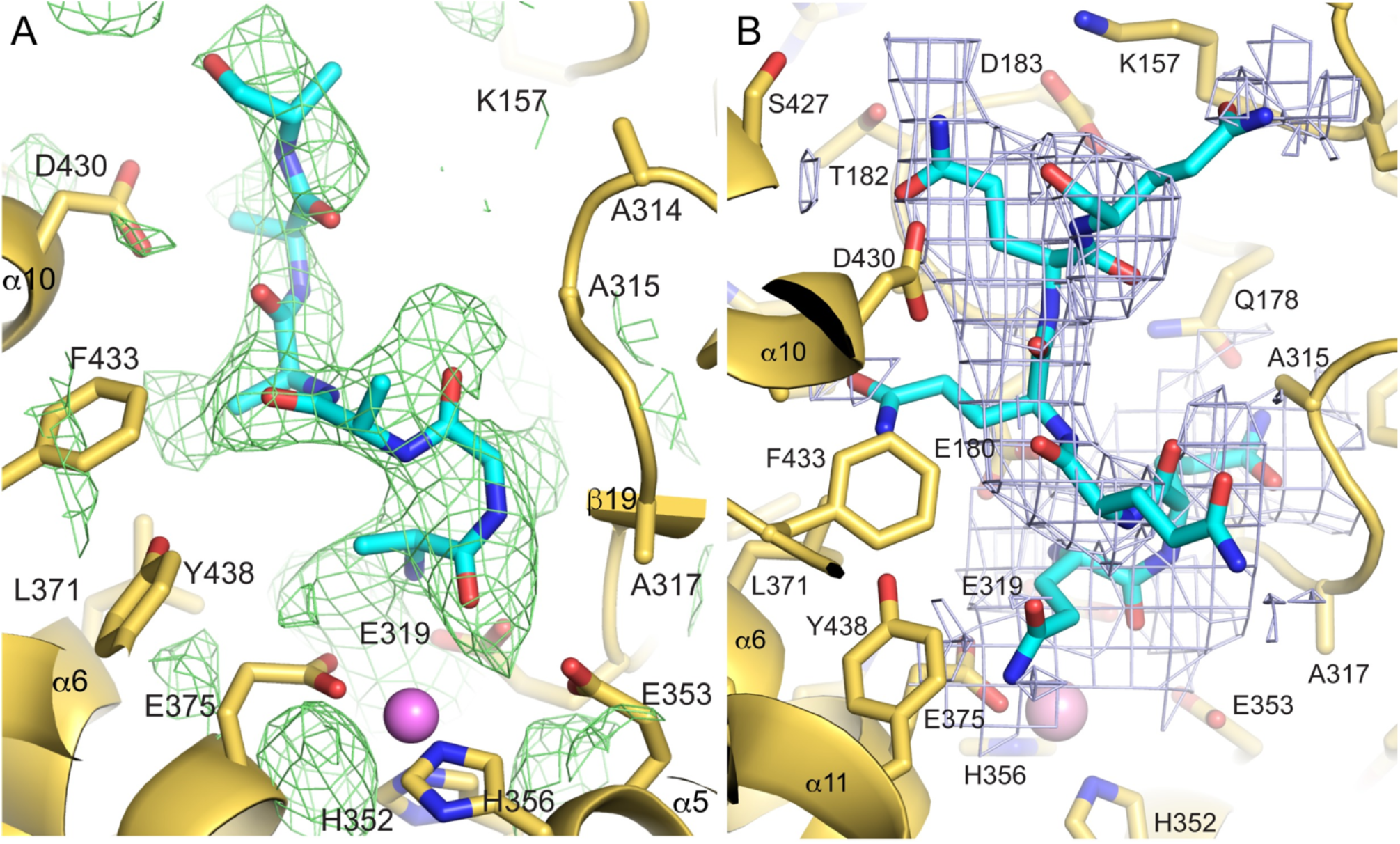
Polyglutamine peptide binding by PSA. (A) Weighted difference electron density (green mesh, 1.5 σ cutoff) in the PSA active site region was calculated with 3.6 Å resolution data obtained from a crystal soaked in a solution containing 200 μM polyglutamine peptide (Lys_2_Gln_15_Lys_2_). A polyalanine peptide (cyan carbons) is shown modeled into the density to illustrate the path of the bound peptide. Nearby elements of the catalytic domain are shown in gold with the side chains of active site and potentially interacting residues shown in stick representation (gold carbons). (B) Refined peptide complex with the bound peptide built as polyglutamine. The peptide is shown with cyan carbons and the protein in gold with all side chains in stick representation. 2Fo-Fc electron density for the peptide is in light blue mesh (0.26 sigma cutoff). The active site zinc ion is shown as a pink sphere in both panels.

No major changes in the overall conformation of PSA are evident from the electron density upon binding the PQ substrate. Since the peptide was soaked into PSA crystals, it is likely that lattice contacts in the crystal prevent a conformational change in PSA despite the presence of substrate at the active site.

As noted, the interaction with PQ appears to involve Phe433. To further assess the role of this residue, it was mutated to alanine (PSA^F433A^) and the mutant protein produced for kinetic studies in comparison with wild type PSA. K_i_ values were determined for the PQ peptide and a reference substrate, dynorphin A(1-17), by competitive inhibition of the fluorogenic substrate alanine 4-methoxy-β-naphthylamide (Ala-4MβNA) (**Fig 10**, data for graphs in **S1 Table**). Both PQ and dynorphin A(1-17) were found to be competitive with the fluorogenic substrates (**Fig 10A,B**). The apparent K_i_ for the PQ peptide with wild type PSA was 1.3 µM, 95% CI [1.10-1.54] (**Table 2**). The K_i_ of PQ with the F433A mutant was found to be 4.9 µM, 95% CI [2.9-7.9], or 3.6 fold higher than wild type. Thus, mutating Phe433 reduces affinity for the PQ peptide, consistent with it playing a role in binding as indicated by the crystal structure. Interestingly, mutating Phe433 also decreased affinity for the reference peptide, dynorphin A(1-17). The K_i_ with PSA^F433A^, 2.6 μM, 95% CI [2.38-2.82], was 5.9 fold higher than the K_i_ with wild type PSA, 0.44 μM, 95% CI [0.38-0.51]. Either dynorphin A(1-17) interacts in a similar manner as PQ or mutating F433 has a more general effect on substrate binding.

**Figure 10.**
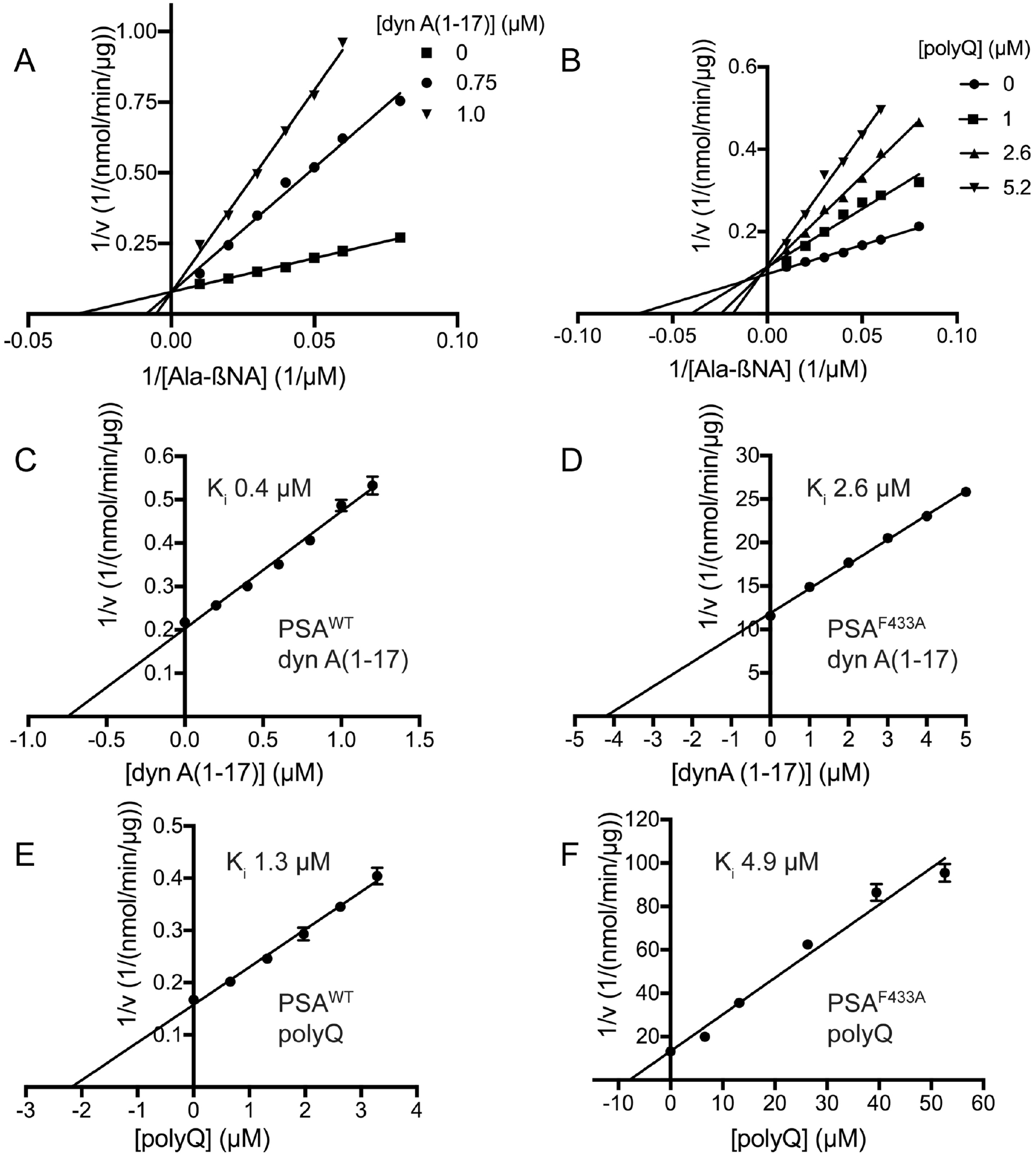
Interaction with polyglutamine peptide (PQ) and dynorphin A(1-17). Binding of the two peptides was followed by their ability to inhibit hydrolysis of a standard fluorogenic peptide substrate, alanine 4-methoxy-β-naphthylamide. Double reciprocal plots for inhibition by (A) dyn A(1-17) and (B) PQ indicate the peptides act as competitive inhibitors of the fluorogenic substrate. Single reciprocal plots (Dixon plots) for dyn A(1-17) interacting with (C) PSA^wt^ and (D) PSA^F433A^ are shown along with PQ interacting with (E) PSA^wt^ and (F) PSA^F433A^. In these plots, the abscissa intercept is –K_i_(1+[S]/K_m_), where [S] is the concentration and K_m_ the Michaelis constant of the fluorogenic substrate. Error bars represent standard error of the mean for triplicate measurements.

**Table 2.** Apparent K_i_ values for polyglutamine and dynorphin A(1-17) with PSA^wt^ and PSA^F433A^.

| enzyme | $K_i$ | |
| --- | --- | --- |
| | polyglutamine (μM) | dynorphin A(1-17) (μM) |
| PSA<sup>wt</sup> | 1.30 (1.10-1.54)<sup>a</sup> | 0.44 (0.38-0.51) |
| PSA<sup>F433A</sup> | 4.9 (2.9-7.2) | 2.60 (2.38-2.82) |

<sup>a</sup> 95% confidence interval

## Discussion

Most of the cytosolic aminopeptidase activity in the mammalian brain and likely other tissues is attributable to puromycin sensitive aminopeptidase (PSA). It was first purified from rat brain [96, 97] and bovine brain [8] based on its cleavage of enkephalin. PSA orthologs have been found in a wide range of organisms, including plants [98], primitive eukaryotes [22, 23] and amphibians [99], and the presence of clear orthologs across Eukarya suggests an essential function for PSA. M1 aminopeptidases exist in Archaea [100] and bacteria [101], indicating that the family is of ancient origin. The work reported here shows a close structural similarity between PSA and these distantly related prokaryotic enzymes.

As noted, *in vitro* and *in vivo* studies have identified a number of substrates for PSA [9, 102, 103] [8, 104]. A key feature of PSA, therefore, is its ability to accommodate a number of substrate amino acid sequences in its active site. It is clear that, although preferences exist, various types of amino acids can be accommodated at any position relative to the cleavage site. While peptidases frequently do not have absolute specificities at particular positions, the ability to recognize such a broad range of seemingly unrelated sequences is often a characteristic of zinc metallopeptidases that metabolize bioactive peptides [3, 105–108].

The PSA structure suggests two factors that may contribute to the broad substrate specificity of the enzyme. In the related APN with bound bestatin [43], the position of the phenyl group of bestatin likely defines the S1 subsite of the enzyme. Interestingly, Met260, which forms part of the subsite, must change conformation in order to accommodate the bulky phenyl group [44]. LTA4H has an even larger tyrosine residue at the equivalent position [46]. On the other hand, PSA has a much smaller alanine residue (Ala315) at this site. The smaller residue in PSA opens up the site relative to the other aminopeptidases, suggesting that it may be even less selective at the P1 substrate position. In the APN complex, a glutamine residue is also present in the S1 subsite, and this residue is conserved in PSA (Gln178) and LTA4H. In factor F3, however, this residue is a histidine (His99) [38]. FactorF3 prefers negatively charged residues at the P1 position, and the substitution of the at least partially positive histidine for the polar glutamine at this position likely accounts for that preference. The nature of the residues in the likely S1 subsite of PSA, particularly the presence of the small Ala315 and the polar Gln178, may allow for a broad range of residues at substrate P1 position. In addition to these considerations at P1, the presence of many aromatic and hydrophobic residues around the putative S1 subsite and other regions near the active site may mediate broad specificity by allowing different substrates to interact with different portions of this flat, carbon-rich surface.

The effect on PQ binding in the F433A mutant serves to support that the electron density seen in the complex crystal structure does reflect the backbone path of the PQ peptide. In addition, the peptide acting as a competitive inhibitor of a small fluorogenic substrate is consistent with the catalytic binding mode observed in the crystal structure. Phenylalanine is generally conserved at the equivalent of position 433 in other M1 aminopeptidases, except in PfA-M1 and LTA4H where there is a conservative change to tyrosine. Structures of peptide analog inhibitors or peptides complexed with M1 family aminopeptidases align well in the active site, but the paths of the ligand backbones diverge as they extend toward what would be the C termini of bound substrates (**Fig 11A**). Interestingly, bestatin bound to APN extends in the direction of the PQ peptide bound to PSA, although the peptide mimic bestatin is in the opposite orientation, with its C-terminal carboxyl group coordinating the active site zinc ion. This, diversity of interactions, taken with the binding path of PQ reported here, supports the proposal that the surface near the active site can accommodate a number of substrate binding modes. Additional crystal structures of PSA with different bound peptide substrates will be needed to test this proposal. Since PSA in the crystals described here is likely constrained by lattice contacts to remain in the open conformation, ideally additional structures would be determined with pre-formed enzyme-substrate complexes to enhance relevance to interaction in solution.

**Figure 11.**
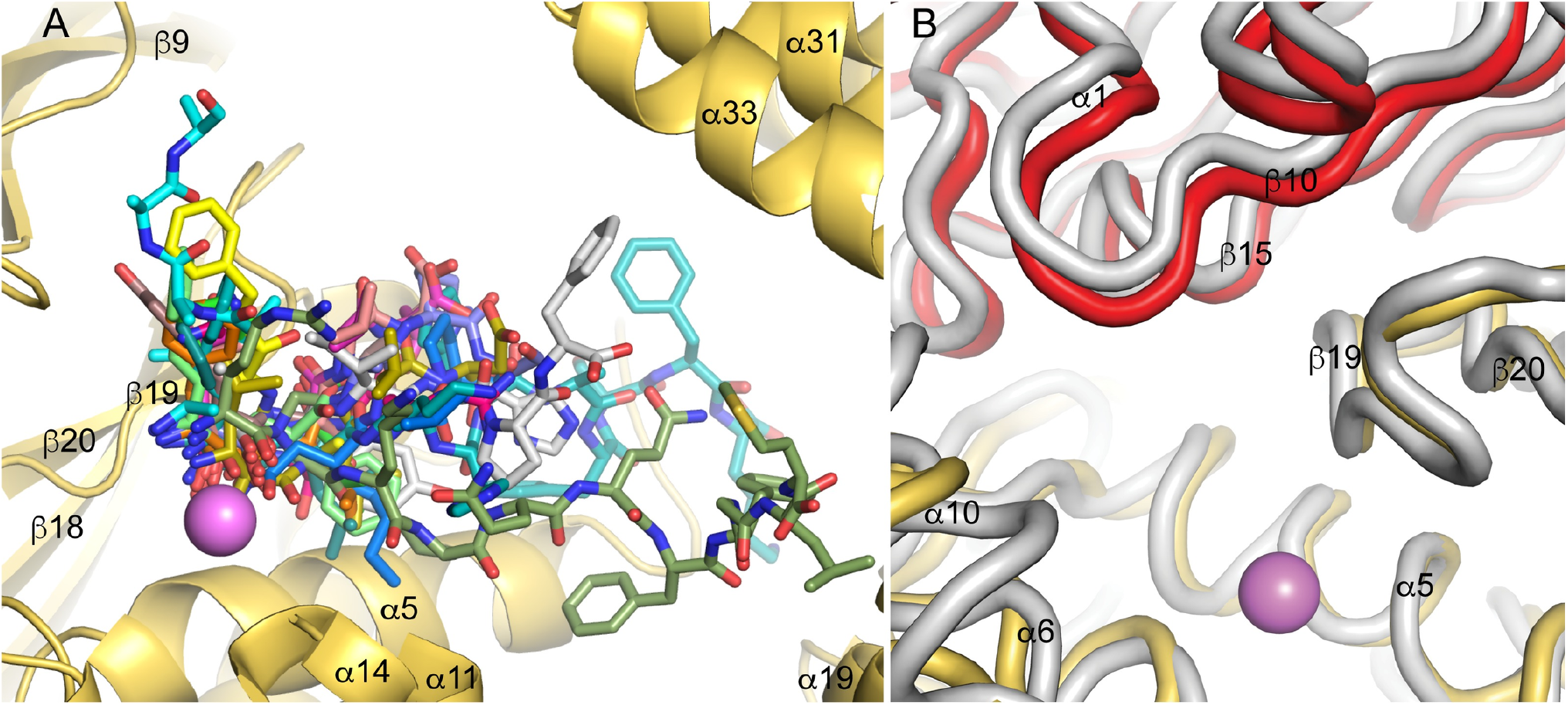
Substrate interactions. (A) Positions of substrates and substrate analogs over the binding surfaces of M1 family peptidases. Selected peptides or peptide analogs from M1 aminopeptidase structures superimposed on PSA are shown in stick format with different carbon colors. The active site zinc ion is shown as a pink sphere and elements of the catalytic domain are in gold. The polyalanine model built into the PSA-polyglutamine structure is shown with carbons colored in cyan. Other structures shown are APA with bestatin [74] (PDB ID 4KXB, green carbons), APA with amastatin [74] (4KX8, magenta), APN with bestatin [75] (4FYR, yellow), APN with amastatin [75] (4FYT, salmon), APN with angiotensin IV [75] (4FYS, gray), AnAPN1 with 5-mer peptide [78] (4WZ9, slate), PfA-M1 with bestatin [45] (3EBH, orange), PfA-M1 with phosphinic dipeptide analog [45] (3EBI, lime), LTA4H with bestatin [46] (1HS6, dark teal), LTA4H with a 3-mer peptide [109] (3B7S, hot pink), ePepN with actinonin [110] (4Q4E, marine), ePepN with amastatin [110] (4Q4I, olive), porcine APN with substance P [76] (4HOM, split pea), ERAP1 with a 10-mer phosphinic peptide [59] (6RQX, teal), and M1dr with a 3-mer peptide [84] (6IFG, dark violet). (B) Hinge motion at the interface between PSA domains 1 and 2. Cα traces of domain 1 (red) and domain 2 (gold) from the crystal structure are shown superimposed on the trace of a structure from normal mode analysis (gray) using the NOMAD-Ref server [111]. Movement of domain 1 in the normal mode analysis relative to its position in the crystal structure can be seen as a shift of the gray trace toward the top of the figure.

The M1 family peptidases have been crystallized in two overall conformations differing by a hinge-like motion of the C-terminal domain IV relative to the long, N-terminal arm (domains I-III) of the V-shaped enzyme. In the majority of the crystal structures, the C-terminal domain is closed over the active site, interacting extensively with domain II, which restricts access and possible substrate binding modes. In contrast, PSA, Factor F3, and forms of ERPA1 and APN adopt open conformations, with domain IV rotated away from the N-terminal arm by about 40° in most cases. It has not, however, been established whether all the M1 peptidases sample both open and closed conformations in solution (with perhaps different equilibrium distributions for the different peptidases). The observations that ERAP1 crystallizes in both conformations [47, 48], and that different Factor F3 molecules in the crystal asymmetric unit show different rotations of domain IV [38] suggest that in at least in some cases the relative positions of the N- and C-terminal arms can vary dynamically. The consequences of this conformational dynamics for the range of substrate binding modes remain to be established.

Addlagatta and colleagues have suggested a binding mode for the PSA specific inhibitor puromycin based on the structure of puromycin bound to an inactive mutant of ePepN and docking to a closed form PSA homology model [112]. The nucleoside portion of the inhibitor interacts near the active site zinc ion and coordinating residues, while the remainder of the molecule extends toward helix 31 of domain IV. The open conformation PSA structure reported here was superimposed domain-by-domain on the AlphaFold PSA model to generate a model for the closed conformation of the enzyme. The puromycin binding mode suggested previously is largely compatible with this closed model (**Fig. 12A**) and was used as the starting point for docking with ROSIE Ligand_docking [113–115]. Interestingly, the three lowest energy models showed similar positions and orientations for puromycin (see **Fig. 12A**), The docked puromycin ligands adopt more compact conformations and move away from the active site toward the surface of domain IV relative to the puromycin binding mode proposed earlier. In this position, a number of PSA side chains are placed to interact with the ligand, primarily from helix 11 of domain II and helix 31 of domain IV (**Fig. 12B**). Puromycin soaked into crystals of active ePepN showed hydrolysis products in the active site [112]. Superimposing that structure on the closed model of PSA shows that O-methyl-L-tyrosine (OMT) fits well into the S1 subsite of the PSA closed model (**Fig. 12C**). The puromycin aminonucleoside (PAN) fragment, while bound to ePepN in an orientation different from its position during hydrolysis, is also not obstructed by any groups in the closed PSA model. Therefore, the structure affords no obvious reason why puromycin is not hydrolyzed to any great extent by PSA. Its functioning as a competitive inhibitor likely results from an unproductive, high affinity binding mode, like the one suggested by the docking study, that sterically restricts access to the active site.

**Figure 12.**
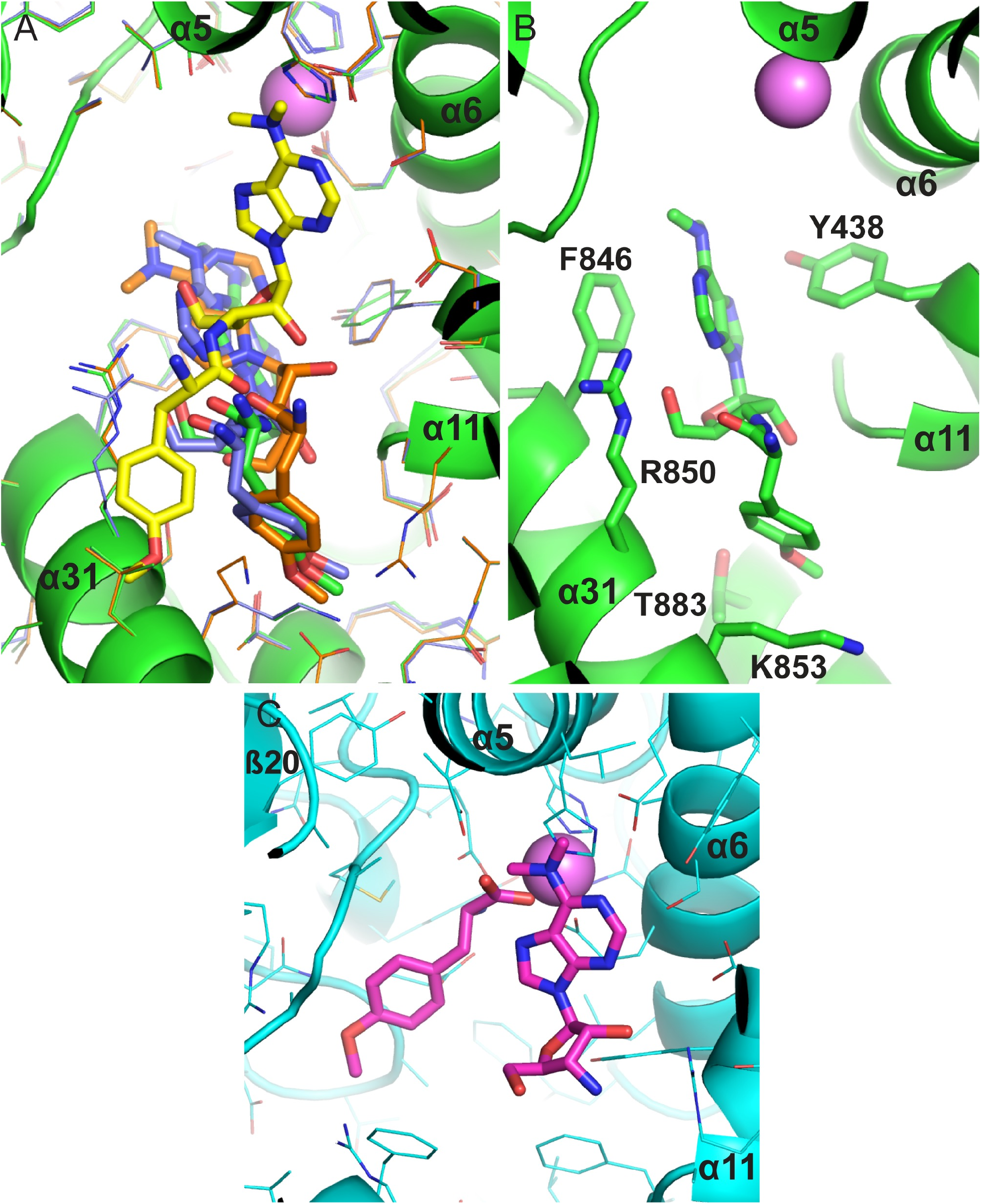
Modeling puromycin interaction with a closed form of PSA. The closed form model for PSA was constructed by superimposing the open PSA crystal structure reported here on the closed form AlphaFold model domain by domain. (A) Docking of puromycin with the closed form PSA using ROSIE Ligand_docking [113–115]. Puromycin shown with yellow carbon atoms is from superposition of an inactive ePepN-puromycin complex reported by Addlagatta and colleagues [112] on the closed PSA model. That puromycin pose was used as the starting point for docking with the closed PSA model. The three lowest energy complexes are show with green, purple, orange carbons, corresponding protein side chains, and the lowest energy model backbone in green. The zinc ion is show as a pink sphere. (B) The lowest energy docked PSA-puromycin model. Side chains positioned to possibly interact with the ligand are shown. (C) Hydrolysis fragments of puromycin superimposed on the closed PSA model. The fragments from the active ePepN-puromycin crystal structure [112] are shown with magenta carbons based on the superposition of the ePepN complex on the closed PSA model. Side chains from the PSA model are shown.

PSA has been implicated in the metabolism of two proteins associated with protein aggregation disorders, Tau and superoxide dismutase [26–29, 33]. In both cases, reports suggest that PSA may play a direct role in degrading these large substrates, and evidence has been presented for endopeptidase activity of PSA. However, a study using purified PSA and Tau failed to find a direct role of PSA in Tau degradation [30]. PSA has been reported to stimulate autophagy [32], and this may at least in part account for its effects on levels of Tau and superoxide dismutase, both of which have been shown to be degraded via macroautophagy as well as other mechanisms [116, 117]. Alternatively, since PSA contains microtubule-binding sequences and has been shown to co-localize with tubulin [11, 62, 118], it is possible that it may influence Tau lifetime by increasing the proportion of protein not bound to microtubules. Localization to microtubules may also play a role in the function of PSA in meiotic cell division, where its absence causes defects in chromosome segregation, recombination, development of cell polarity, and cell cycle progression [22, 98]. Here PSA peptidase activity may be required, since inhibitors reproduce at least some of the effects of gene knockouts.

Despite questions regarding direct degradation of Tau or SOD, it is useful to examine the PSA structure with regard to its potential activity on large substrates or at least interaction with proteins. The open conformation of PSA may allow loop segments from folded or partially folded proteins to enter the active site groove. The structural barrier at one end of the active site, however, makes it unlikely that a loop segment could bind in a productive manner. Since this barrier is largely composed of elements from the N-terminal domain of PSA, one possible mechanism for endolytic cleavage would be the N-terminal domain of PSA swinging away from the metallopeptidase active site region in a hinge like motion (**Fig 11B**). Such a movement of the N-terminal domain would open the closed end of the active site channel, allowing a protein loop segment to extend on either side of the active site for endolytic cleavage. Only a single backbone connection exists between the N-terminal and catalytic domains, and this connecting segment is in an open coil conformation. Therefore, this region might act as the hinge. In this model, the hinge-like conformational change would be a relatively rare event, consistent with the reported poor efficiency of large substrate degradation by PSA [27]. The interface between the N-terminal and catalytic domain is not predominantly hydrophobic, suggesting that exposing the surfaces would not be prohibitively unfavorable. In fact, the largest hydrophobic region at the interface is near the hinge region between the N-terminal and catalytic domains where it would not be greatly exposed by a hinge motion.

In conclusion, the work reported here demonstrates the basis for aminopeptidase activity by PSA and suggests a mechanism for its broad substrate recognition. In addition, the path of polyglutamine substrates is defined, suggesting they may bind in a manner distinct from other peptide substrates.

## Acknowledgements

We thank the staff of Advanced Photon Source beamline 22-ID (SER-CAT) for their help with data collection. Use of the Advanced Photon Source is supported by the U.S. Department of Energy. We also thank Trevor Creamer and Veronique Chellgren for supplying polyglutamine peptide and assistance with its use in this work.

## Supporting information

**Table S1:** Data for graphs in Fig. 10

| Data for Fig. 1A, Ala-βNA series at three dyn A(1-17) concentrations |  |  |  |  |
| --- | --- | --- | --- | --- |
| 1/[Ala-βNA] (1/μM) | 1/v (1/nmol/min/ug) |  |  |  |
|  | 0.0 μM<br>dyn A(1-17) | 0.75 μM<br>dyn A(1-17) | 1.0 μM<br>dyn A(1-17) |  |
| 0.01 | 0.107 | 0.144 | 0.244 |  |
| 0.02 | 0.126 | 0.244 | 0.348 |  |
| 0.03 | 0.150 | 0.348 | 0.495 |  |
| 0.04 | 0.166 | 0.466 | 0.647 |  |
| 0.05 | 0.199 | 0.520 | 0.773 |  |
| 0.06 | 0.223 | 0.621 | 0.960 |  |
| 0.08 | 0.271 | 0.755 |  |  |
| Data for Fig. 1B, Ala-βNA series at four polyQ peptide concentrations |  |  |  |  |
| 1/[Ala-βNA] (1/μM) | 1/v (1/nmol/min/ug) |  |  |  |
|  | 0.0 μM<br>polyQ | 1.0 μM<br>polyQ | 2.6 μM<br>polyQ | 5.2 μM<br>polyQ |
| 0.01 | 0.114 | 0.129 | 0.156 | 0.170 |
| 0.02 | 0.126 | 0.165 | 0.197 | 0.240 |
| 0.03 | 0.137 | 0.199 | 0.254 | 0.337 |
| 0.04 | 0.149 | 0.242 | 0.283 | 0.368 |
| 0.05 | 0.167 | 0.271 | 0.330 | 0.434 |
| 0.06 | 0.180 | 0.288 | 0.391 | 0.495 |
| 0.08 | 0.213 | 0.320 | 0.466 |  |
| Data for Fig. 1C, dyn A(1-17) inhibition of PSA<sup>WT</sup> |  |  |  |  |
| [dyn A(1-17)] (μM) | 1/v (1/nmol/min/ug) |  |  |  |
| 0.0 | 0.216 | 0.217 | 0.219 |  |
| 0.2 | 0.258 | 0.261 | 0.250 |  |
| 0.4 | 0.296 | 0.310 | 0.297 |  |
| 0.6 | 0.343 | 0.364 | 0.345 |  |
| 0.8 | 0.417 | 0.397 | 0.405 |  |
| 1.0 | 0.461 | 0.502 | 0.497 |  |
| 1.2 | 0.574 | 0.517 | 0.508 |  |

| Data for Fig. 1D, dyn A(1-17) inhibition of PSA<sup>F433A</sup> |  |  |  |
| --- | --- | --- | --- |
| [dyn A(1-17)] (μM) | 1/v (1/nmol/min/ug) |  |  |
| 0.0 | 12.094 | 11.530 | 11.031 |
| 1.0 | 15.072 | 14.917 | 14.691 |
| 2.0 | 17.807 | 18.247 | 17.001 |
| 3.0 | 20.352 | 20.438 | 20.745 |
| 4.0 | 23.540 | 22.647 | 22.933 |
| 5.0 | 26.765 | 25.631 | 25.100 |
| Data for Fig. 1E, polyQ inhibition of PSA<sup>WT</sup> |  |  |  |
| [polyQ] (μM) | 1/v (1/nmol/min/ug) |  |  |
| 0.0 | 0.166 | 0.169 | 0.167 |
| 0.7 | 0.204 | 0.196 | 0.206 |
| 1.3 | 0.242 | 0.245 | 0.251 |
| 2.0 | 0.269 | 0.303 | 0.307 |
| 2.6 | 0.334 | 0.345 | 0.356 |
| 3.3 | 0.383 | 0.394 | 0.435 |
| Data for Fig. 1F, polyQ inhibition of PSA<sup>F433A</sup> |  |  |  |
| [polyQ] (μM) | 1/v (1/nmol/min/ug) |  |  |
| 0.0 | 12.667 | 13.681 | 12.903 |
| 6.6 | 18.994 | 20.089 | 20.799 |
| 13.2 | 34.831 | 35.892 | 36.130 |
| 26.3 | 62.515 | 62.605 | 62.247 |
| 39.5 | 83.956 | 93.908 | 81.445 |
| 52.6 | 90.401 | 103.746 | 92.121 |

## References

1. Littlewood GM, Iversen LL, Turner AJ. Neuropeptides and their peptidases: functional considerations. Neurochem Int. 1988;12:383–9.

2. Turner AJ, ed. Neuropeptides and their peptidases.: Ellis Horwood, New York; 1987.

3. Konkoy CS, Davis TP. Ectoenzymes as sites of peptide regulation. Trends in Pharmaceutical Sciences. 1996;17:288–94.

4. McKelvy JF, Blumberg S. Inactivation and metabolism of neuropeptides. Annual Review of Neuroscience. 1986;9:415–34.

5. Rawlings ND, Barrett AJ. Evolutionary families of metallopeptidases. Methods in Enzymology. 1995;248:183–228.

6. Albiston AL, Ye S, Chai SY. Membrane bound members of the M1 family: more than aminopeptidases. Protein Pept Lett. 2004;11(5):491–500.

7. Mantle D. Comparison of soluble aminopeptidases in human cerebral cortex, skeletal muscle and kidney tissues. Clin Chim Acta. 1992;207(1-2):107–18.

8. Hersh LB, McKelvy JF. An aminopeptidase from bovine brain which catalyzes the hydrolysis of enkephalin. J Neurochem. 1981;36(1):171–8.

9. Hersh LB, Smith TE, McKelvy JF. Cleavage of endorphins to des-Tyr endorphins by homogeneous bovine brain aminopeptidase. Nature. 1980;286(5769):160–2.

10. Safavi A, Hersh LB. Degradation of dynorphin-related peptides by the puromycin-sensitive aminopeptidase and aminopeptidase M. J Neurochem. 1995;65(1):389–95.

11. Constam DB, Tobler AR, Rensing-Ehl A, Kemler I, Hersh LB, Fontana A. Puromycin-sensitive aminopeptidase. Sequence analysis, expression, and functional characterization. Journal of Biologial Chemistry. 1995;270(45):26931–9.

12. Peer WA. The role of multifunctional M1 metallopeptidases in cell cycle progression. Ann Bot. 2011;107(7):1171–81.

13. Fortin SM, Marshall SL, Jaeger EC, Greene PE, Brady LK, Isaac RE, et al. The PAM-1 aminopeptidase regulates centrosome positioning to ensure anterior-posterior axis specification in one-cell C. elegans embryos. Dev Biol. 2010;344(2):992–1000.

14. Saturno DM, Castanzo DT, Williams M, Parikh DA, Jaeger EC, Lyczak R. Sustained centrosome-cortical contact ensures robust polarization of the one-cell C. elegans embryo. Dev Biol. 2017;422(2):135–45.

15. Osana S, Kitajima Y, Suzuki N, Nunomiya A, Takada H, Kubota T, et al. Puromycin-sensitive aminopeptidase is required for C2C12 myoblast proliferation and differentiation. J Cell Physiol. 2021;236(7):5293–305.

16. Benton D, Jaeger EC, Kilner A, Kimble A, Lowry J, Schleicher EM, et al. Interactions between the WEE-1.3 kinase and the PAM-1 aminopeptidase in oocyte maturation and the early C. elegans embryo. G3 (Bethesda). 2021;11(4).

17. Agrawal N, Brown MA. Genetic associations and functional characterization of M1 aminopeptidases and immune-mediated diseases. Genes Immun. 2014;15(8):521–7.

18. Lowenberg B, Morgan G, Ossenkoppele GJ, Burnett AK, Zachee P, Duhrsen U, et al. Phase I/II clinical study of Tosedostat, an inhibitor of aminopeptidases, in patients with acute myeloid leukemia and myelodysplasia. J Clin Oncol. 2010;28(28):4333–8.

19. Singh R, Williams J, Vince R. Puromycin based inhibitors of aminopeptidases for the potential treatment of hematologic malignancies. Eur J Med Chem. 2017;139:325–36.

20. Osada T, Watanabe G, Kondo S, Toyoda M, Sakaki Y, Takeuchi T. Male reproductive defects caused by puromycin-sensitive aminopeptidase deficiency in mice. Mol Endocrinol. 2001;15(6):960–71.

21. Osada T, Watanabe G, Sakaki Y, Takeuchi T. Puromycin-sensitive aminopeptidase is essential for the maternal recognition of pregnancy in mice. Mol Endocrinol. 2001;15(6):882–93.

22. Lyczak R, Zweier L, Group T, Murrow MA, Snyder C, Kulovitz L, et al. The puromycin-sensitive aminopeptidase PAM-1 is required for meiotic exit and anteroposterior polarity in the one-cell Caenorhabditis elegans embryo. Development. 2006;133(21):4281–92.

23. Brooks DR, Hooper NM, Isaac RE. The Caenorhabditis elegans orthologue of mammalian puromycin-sensitive aminopeptidase has roles in embryogenesis and reproduction. Journal of Biologial Chemistry. 2003;278(44):42795–801.

24. Schulz C, Perezgasga L, Fuller MT. Genetic analysis of dPsa, the Drosophila orthologue of puromycin-sensitive aminopeptidase, suggests redundancy of aminopeptidases. Dev Genes Evol. 2001;211(12):581–8.

25. Osada T, Ikegami S, Takiguchi-Hayashi K, Yamazaki Y, Katoh-Fukui Y, Higashinakagawa T, et al. Increased anxiety and impaired pain response in puromycin-sensitive aminopeptidase gene-deficient mice obtained by a mouse gene-trap method. Journal of Neuroscience. 1999;19(14):6068–78.

26. Karsten SL, Sang TK, Gehman LT, Chatterjee S, Liu J, Lawless GM, et al. A genomic screen for modifiers of tauopathy identifies puromycin-sensitive aminopeptidase as an inhibitor of tau-induced neurodegeneration. Neuron. 2006;51(5):549–60.

27. Sengupta S, Horowitz PM, Karsten SL, Jackson GR, Geschwind DH, Fu Y, et al. Degradation of tau protein by puromycin-sensitive aminopeptidase in vitro. Biochemistry. 2006;45(50):15111–9.

28. Yanagi K, Tanaka T, Kato K, Sadik G, Morihara T, Kudo T, et al. Involvement of puromycin-sensitive aminopeptidase in proteolysis of tau protein in cultured cells, and attenuated proteolysis of frontotemporal dementia and parkinsonism linked to chromosome 17 (FTDP-17) mutant tau. Psychogeriatrics. 2009;9(4):157–66.

29. Kudo LC, Parfenova L, Ren G, Vi N, Hui M, Ma Z, et al. Puromycin-sensitive aminopeptidase (PSA/NPEPPS) impedes development of neuropathology in hPSA/TAU(P301L) double-transgenic mice. Hum Mol Genet. 2011;20(9):1820–33.

30. Chow KM, Guan H, Hersh LB. Aminopeptidases do not directly degrade tau protein. Mol Neurodegener. 2010;5:48.

31. Bhutani N, Venkatraman P, Goldberg AL. Puromycin-sensitive aminopeptidase is the major peptidase responsible for digesting polyglutamine sequences released by proteasomes during protein degradation. EMBO Journal. 2007;26(5):1385–96.

32. Menzies FM, Hourez R, Imarisio S, Raspe M, Sadiq O, Chandraratna D, et al. Puromycin-sensitive aminopeptidase protects against aggregation-prone proteins via autophagy. Hum Mol Genet. 2010;19(23):4573–86.

33. Ren G, Ma Z, Hui M, Kudo LC, Hui KS, Karsten SL. Cu, Zn-superoxide dismutase 1 (SOD1) is a novel target of Puromycin-sensitive aminopeptidase (PSA/NPEPPS): PSA/NPEPPS is a possible modifier of amyotrophic lateral sclerosis. Mol Neurodegener. 2011;6:29.

34. Thompson MW, Govindaswami M, Hersh LB. Mutation of active site residues of the puromycin-sensitive aminopeptidase: conversion of the enzyme into a catalytically inactive binding protein. Arch, Biochem, Biophys,. 2003;413(2):236–42.

35. Thompson MW, Hersh LB. Analysis of conserved residues of the human puromycin-sensitive aminopeptidase. Peptides. 2003;24(9):1359–65.

36. Rodgers DW. Practical Cryocrystallography. Methods Enzymol. 1997;276:183–203.

37. Otwinowski Z, Minor W. Processing of X-ray diffraction data collected in oscillation mode. Macromolecular Crystallography, Pt A. 1997;276:307–26.

38. Kyrieleis OJ, Goettig P, Kiefersauer R, Huber R, Brandstetter H. Crystal structures of the tricorn interacting factor F3 from Thermoplasma acidophilum, a zinc aminopeptidase in three different conformations. Journal of Molecular Biology. 2005;349(4):787–800.

39. Liebschner D, Afonine PV, Baker ML, Bunkoczi G, Chen VB, Croll TI, et al. Macromolecular structure determination using X-rays, neutrons and electrons: recent developments in Phenix. Acta Crystallogr D Struct Biol. 2019;75(Pt 10):861–77.

40. Adams PD, Afonine PV, Bunkoczi G, Chen VB, Davis IW, Echols N, et al. PHENIX: a comprehensive Python-based system for macromolecular structure solution. Acta Crystallogr D Biol Crystallogr. 2010;66(Pt 2):213–21.

41. Emsley P, Cowtan K. Coot: model-building tools for molecular graphics. Acta Crystallogr D Biol Crystallogr. 2004;60(Pt 12 Pt 1):2126–32.

42. Chen S, Wetzel R. Solubilization and disaggregation of polyglutamine peptides. Protein Science. 2001;10(4):887–91.

43. Addlagatta A, Gay L, Matthews BW. Structure of aminopeptidase N from Escherichia coli suggests a compartmentalized, gated active site. Proc Natl Acad Sci U S A. 2006;103(36):13339–44.

44. Ito K, Nakajima Y, Onohara Y, Takeo M, Nakashima K, Matsubara F, et al. Crystal structure of aminopeptidase N (proteobacteria alanyl aminopeptidase) from Escherichia coli and conformational change of methionine 260 involved in substrate recognition. Journal of Biologial Chemistry. 2006;281(44):33664–76.

45. McGowan S, Porter CJ, Lowther J, Stack CM, Golding SJ, Skinner-Adams TS, et al. Structural basis for the inhibition of the essential Plasmodium falciparum M1 neutral aminopeptidase. Proc Natl Acad Sci U S A. 2009;106(8):2537–42.

46. Thunnissen MM, Nordlund P, Haeggstrom JZ. Crystal structure of human leukotriene A(4) hydrolase, a bifunctional enzyme in inflammation. Nat Struct Biol. 2001;8(2):131–5.

47. Kochan G, Krojer T, Harvey D, Fischer R, Chen L, Vollmar M, et al. Crystal structures of the endoplasmic reticulum aminopeptidase-1 (ERAP1) reveal the molecular basis for N-terminal peptide trimming. Proc Natl Acad Sci U S A. 2011;108(19):7745–50.

48. Nguyen TT, Chang SC, Evnouchidou I, York IA, Zikos C, Rock KL, et al. Structural basis for antigenic peptide precursor processing by the endoplasmic reticulum aminopeptidase ERAP1. Nat Struct Mol Biol. 2011;18(5):604–13.

49. Birtley JR, Saridakis E, Stratikos E, Mavridis IM. The crystal structure of human endoplasmic reticulum aminopeptidase 2 reveals the atomic basis for distinct roles in antigen processing. Biochemistry. 2012;51(1):286–95.

50. Holmes MA, Matthews BW. Structure of thermolysin refined at 1.6 A resolution. Journal of Molecular Biology. 1982;160(4):623–39.

51. Holmes MA, Tronrud DE, Matthews BW. Structural analysis of the inhibition of thermolysin by an active-site-directed irreversible inhibitor. Biochemistry. 1983;22(1):236–40.

52. Colman PM, Jansonius JN, Matthews BW. The structure of thermolysin: an electron density map at 2-3 A resolution. Journal of Molecular Biology. 1972;70(3):701–24.

53. Matthews BW, Colman PM, Jansonius JN, Titani K, Walsh KA, Neurath H. Structure of thermolysin. Nat New Biol. 1972;238(80):41–3.

54. Matthews BW, Jansonius JN, Colman PM, Schoenborn BP, Dupourque D. Three-dimensional structure of thermolysin. Nat New Biol. 1972;238(80):37–41.

55. Matthews BW, Weaver LH, Kester WR. The conformation of thermolysin. Journal of Biologial Chemistry. 1974;249:8030–44.

56. Rawlings ND, Tolle DP, Barrett AJ. MEROPS: the peptidase database. Nucleic Acids Research. 2004;32(Database issue):D160–4.

57. Andrade MA, Petosa C, O’Donoghue SI, Muller CW, Bork P. Comparison of ARM and HEAT protein repeats. Journal of Molecular Biology. 2001;309(1):1–18.

58. Ma Z, Daquin A, Yao J, Rodgers D, Thompson MW, Hersh LB. Proteolytic cleavage of the puromycin-sensitive aminopeptidase generates a substrate binding domain. Archives of Biochemistry and Biophysics. 2003;415(1):80–6.

59. Giastas P, Mpakali A, Papakyriakou A, Lelis A, Kokkala P, Neu M, et al. Mechanism for antigenic peptide selection by endoplasmic reticulum aminopeptidase 1. Proc Natl Acad Sci U S A. 2019;116(52):26709–16.

60. Maben Z, Arya R, Rane D, An WF, Metkar S, Hickey M, et al. Discovery of Selective Inhibitors of Endoplasmic Reticulum Aminopeptidase 1. Journal of Medicinal Chemistry. 2020;63(1):103–21.

61. Maben Z, Arya R, Georgiadis D, Stratikos E, Stern LJ. Conformational dynamics linked to domain closure and substrate binding explain the ERAP1 allosteric regulation mechanism. Nat Commun. 2021;12(1):5302.

62. Kakuta H, Koiso Y, Nagasawa K, Hashimoto Y. Fluorescent bioprobes for visualization of puromycin-sensitive aminopeptidase in living cells. Bioorg Med Chem Lett. 2003;13(1):83–6.

63. Aizawa H, Kawasaki H, Murofushi H, Kotani S, Suzuki K, Sakai H. A common amino acid sequence in 190-kDa microtubule-associated protein and tau for the promotion of microtubule assembly. Journal of Biologial Chemistry. 1989;264(10):5885–90.

64. Lee G, Neve RL, Kosik KS. The microtubule binding domain of tau protein. Neuron. 1989;2(6):1615–24.

65. Al-Bassam J, Ozer RS, Safer D, Halpain S, Milligan RA. MAP2 and tau bind longitudinally along the outer ridges of microtubule protofilaments. J Cell Biol. 2002;157(7):1187–96.

66. Kadavath H, Hofele RV, Biernat J, Kumar S, Tepper K, Urlaub H, et al. Tau stabilizes microtubules by binding at the interface between tubulin heterodimers. Proc Natl Acad Sci U S A. 2015;112(24):7501–756.

67. Dehmelt L, Halpain S. The MAP2/Tau family of microtubule-associated proteins. Genome Biol. 2005;6(1):204.

68. Ravelli RB, Gigant B, Curmi PA, Jourdain I, Lachkar S, Sobel A, et al. Insight into tubulin regulation from a complex with colchicine and a stathmin-like domain. Nature. 2004;428(6979):198–202.

69. Al-Bassam J, Larsen NA, Hyman AA, Harrison SC. Crystal structure of a TOG domain: conserved features of XMAP215/Dis1-family TOG domains and implications for tubulin binding. Structure. 2007;15(3):355–62.

70. Ayaz P, Ye X, Huddleston P, Brautigam CA, Rice LM. A TOG:alphabeta-tubulin complex structure reveals conformation-based mechanisms for a microtubule polymerase. Science. 2012;337(6096):857–60.

71. Leano JB, Rogers SL, Slep KC. A cryptic TOG domain with a distinct architecture underlies CLASP-dependent bipolar spindle formation. Structure. 2013;21(6):939–50.

72. Brouhard GJ, Rice LM. The contribution of alphabeta-tubulin curvature to microtubule dynamics. Journal of Cell Biology. 2014;207(3):323–34.

73. Groll M, Ditzel L, Lowe J, Stock D, Bochtler M, Bartunik HD, et al. Structure of 20S proteasome from yeast at 2.4 A resolution. Nature. 1997;386(6624):463–71.

74. Yang Y, Liu C, Lin YL, Li F. Structural insights into central hypertension regulation by human aminopeptidase A. Journal of Biologial Chemistry. 2013;288(35):25638–45.

75. Wong AH, Zhou D, Rini JM. The X-ray crystal structure of human aminopeptidase N reveals a novel dimer and the basis for peptide processing. Journal of Biologial Chemistry. 2012;287(44):36804–13.

76. Chen L, Lin YL, Peng G, Li F. Structural basis for multifunctional roles of mammalian aminopeptidase N. Proc Natl Acad Sci U S A. 2012;109(44):17966–71.

77. Reguera J, Santiago C, Mudgal G, Ordono D, Enjuanes L, Casasnovas JM. Structural bases of coronavirus attachment to host aminopeptidase N and its inhibition by neutralizing antibodies. PLoS Pathog. 2012;8(8):e1002859.

78. Atkinson SC, Armistead JS, Mathias DK, Sandeu MM, Tao D, Borhani-Dizaji N, et al. The Anopheles-midgut APN1 structure reveals a new malaria transmission-blocking vaccine epitope. Nat Struct Mol Biol. 2015;22(7):532–9.

79. Hermans SJ, Ascher DB, Hancock NC, Holien JK, Michell BJ, Chai SY, et al. Crystal structure of human insulin-regulated aminopeptidase with specificity for cyclic peptides. Protein Science. 2015;24(2):190–9.

80. Mpakali A, Saridakis E, Harlos K, Zhao Y, Papakyriakou A, Kokkala P, et al. Crystal Structure of Insulin-Regulated Aminopeptidase with Bound Substrate Analogue Provides Insight on Antigenic Epitope Precursor Recognition and Processing. J Immunol. 2015;195(6):2842–51.

81. Bauvois C, Jacquamet L, Huston AL, Borel F, Feller G, Ferrer JL. Crystal structure of the cold-active aminopeptidase from Colwellia psychrerythraea, a close structural homologue of the human bifunctional leukotriene A4 hydrolase. Journal of Biologial Chemistry. 2008;283(34):23315–25.

82. Marapaka AK, Pillalamarri V, Gumpena R, Haque N, Bala SC, Jangam A, et al. Discovery, Structural and Biochemical Studies of a rare Glu/Asp Specific M1 Class Aminopeptidase from Legionella pneumophila. Int J Biol Macromol. 2018;120(Pt A):1111–8.

83. Agrawal R, Goyal VD, Singh R, Kumar A, Jamdar SN, Kumar A, et al. Structural basis for the unusual substrate specificity of unique two-domain M1 metallopeptidase. Int J Biol Macromol. 2020;147:304–13.

84. Agrawal R, Goyal VD, Kumar A, Gaur NK, Jamdar SN, Kumar A, et al. Two-domain aminopeptidase of M1 family: Structural features for substrate binding and gating in absence of C-terminal domain. J Struct Biol. 2019;208(1):51–60.

85. Mpakali A, Saridakis E, Harlos K, Zhao Y, Kokkala P, Georgiadis D, et al. Ligand-Induced Conformational Change of Insulin-Regulated Aminopeptidase: Insights on Catalytic Mechanism and Active Site Plasticity. Journal of Medicinal Chemistry. 2017;60(7):2963–72.

86. Jumper J, Evans R, Pritzel A, Green T, Figurnov M, Ronneberger O, et al. Highly accurate protein structure prediction with AlphaFold. Nature. 2021;596(7873):583–9.

87. Varadi M, Anyango S, Deshpande M, Nair S, Natassia C, Yordanova G, et al. AlphaFold Protein Structure Database: massively expanding the structural coverage of protein-sequence space with high-accuracy models. Nucleic Acids Res. 2022;50(D1):D439–D44.

88. Tian W, Chen C, Lei X, Zhao J, Liang J. CASTp 3.0: computed atlas of surface topography of proteins. Nucleic Acids Res. 2018;46(W1):W363–W7.

89. Iturrioz X, Rozenfeld R, Michaud A, Corvol P, Llorens-Cortes C. Study of asparagine 353 in aminopeptidase A: characterization of a novel motif (GXMEN) implicated in exopeptidase specificity of monozinc aminopeptidases. Biochemistry. 2001;40(48):14440–8.

90. Vazeux G, Iturrioz X, Corvol P, Llorens-Cortes C. A glutamate residue contributes to the exopeptidase specificity in aminopeptidase A. Biochemical Journal. 1998;334 (Pt 2):407–13.

91. Cattaneo E, Zuccato C, Tartari M. Normal huntingtin function: an alternative approach to Huntington’s disease. Nat Rev Neurosci. 2005;6(12):919–30.

92. Zoghbi HY, Orr HT. Glutamine repeats and neurodegeneration. Annu Rev Neurosci. 2000;23:217–47.

93. Scherzinger E, Sittler A, Schweiger K, Heiser V, Lurz R, Hasenbank R, et al. Self-assembly of polyglutamine-containing huntingtin fragments into amyloid-like fibrils: implications for Huntington’s disease pathology. Proc Natl Acad Sci U S A. 1999;96(8):4604–9.

94. Davies SW, Turmaine M, Cozens BA, DiFiglia M, Sharp AH, Ross CA, et al. Formation of neuronal intranuclear inclusions underlies the neurological dysfunction in mice transgenic for the HD mutation. Cell. 1997;90(3):537–48.

95. Venkatraman P, Wetzel R, Tanaka M, Nukina N, Goldberg AL. Eukaryotic proteasomes cannot digest polyglutamine sequences and release them during degradation of polyglutamine-containing proteins. Mol Cell. 2004;14(1):95–104.

96. Schnebli HP, Phillipps MA, Barclay RK. Isolation and characterization of an enkephalin-degrading aminopeptidase from rat brain. Biochemica Biophysica Acta. 1979;569(1):89–98.

97. Wagner GW, Tavianini MA, Herrmann KM, Dixon JE. Purification and characterization of an enkephalin aminopeptidase from rat brain. Biochemistry. 1981;20(13):3884–90.

98. Sanchez-Moran E, Jones GH, Franklin FC, Santos JL. A puromycin-sensitive aminopeptidase is essential for meiosis in Arabidopsis thaliana. Plant Cell. 2004;16(11):2895–909.

99. Klein SL, Strausberg RL, Wagner L, Pontius J, Clifton SW, Richardson P. Genetic and genomic tools for Xenopus research: The NIH Xenopus initiative. Dev Dyn. 2002;225(4):384–91.

100. Tamura N, Lottspeich F, Baumeister W, Tamura T. The role of tricorn protease and its aminopeptidase-interacting factors in cellular protein degradation. Cell. 1998;95(5):637–48.

101. Chandu D, Kumar A, Nandi D. PepN, the major Suc-LLVY-AMC-hydrolyzing enzyme in Escherichia coli, displays functional similarity with downstream processing enzymes in Archaea and eukarya. Implications in cytosolic protein degradation. Journal of Biologial Chemistry. 2003;278(8):5548–56.

102. Hui KS, Hui MP, Lajtha A. Major rat brain membrane-associated and cytosolic enkephalin-degrading aminopeptidases: comparison studies. J Neurosci Res. 1988;20(2):231–40.

103. McDermott JR, Mantle D, Lauffart B, Kidd AM. Purification and characterization of a neuropeptide-degrading aminopeptidase from human brain. J Neurochem. 1985;45(3):752–9.

104. McLellan S, Dyer SH, Rodriguez G, Hersh LB. Studies on the tissue distribution of the puromycin-sensitive enkephalin-degrading aminopeptidases. J Neurochem. 1988;51(5):1552–9.

105. Camargo AC, Gomes MD, Toffoletto O, Ribeiro MJ, Ferro ES, Fernandes BL, et al. Structural requirements of bioactive peptides for interaction with endopeptidase 22.19. Neuropeptides. 1994;26(4):281–7.

106. Camargo ACM, Gomes MD, Reichl AP, Ferro ES, Jacchieri S, Hirata IY, et al. Structural features that make oligopeptides susceptible substrates for hydrolysis by recombinant thimet oligopeptidase. Biochemical Journal. 1997;324:517–22.

107. Csuhai E, Safavi A, Thompson MW, Hersh LB. Proteolytic inactivation of secreted neuropeptides. In: Hook V, editor. Proteolytic and Cellular Mechanisms in Prohormone and Neuropeptide Precursor Processing. Heidelberg: Springer-Verlag; 1998.

108. Oliveira V, Campos M, Melo RL, Ferro ES, Camargo AC, Juliano MA, et al. Substrate specificity characterization of recombinant metallo oligo-peptidases thimet oligopeptidase and neurolysin. Biochemistry. 2001;40(14):4417–25.

109. Tholander F, Muroya A, Roques BP, Fournie-Zaluski MC, Thunnissen MM, Haeggstrom JZ. Structure-based dissection of the active site chemistry of leukotriene A4 hydrolase: implications for M1 aminopeptidases and inhibitor design. Chem Biol. 2008;15(9):920–9.

110. Ganji RJ, Reddi R, Gumpena R, Marapaka AK, Arya T, Sankoju P, et al. Structural basis for the inhibition of M1 family aminopeptidases by the natural product actinonin: Crystal structure in complex with E. coli aminopeptidase N. Protein Science. 2015;24(5):823–31.

111. Lindahl E, Azuara C, Koehl P, Delarue M. NOMAD-Ref: visualization, deformation and refinement of macromolecular structures based on all-atom normal mode analysis. Nucleic Acids Research. 2006;34(Web Server issue):W52–6.

112. Reddi R, Ganji RJ, Marapaka AK, Bala SC, Yerra NV, Haque N, et al. Puromycin, a selective inhibitor of PSA acts as a substrate for other M1 family aminopeptidases: Biochemical and structural basis. Int J Biol Macromol. 2020;165(Pt A):1373–81.

113. Combs SA, Deluca SL, Deluca SH, Lemmon GH, Nannemann DP, Nguyen ED, et al. Small-molecule ligand docking into comparative models with Rosetta. Nature protocols. 2013;8(7):1277–98.

114. DeLuca S, Khar K, Meiler J. Fully Flexible Docking of Medium Sized Ligand Libraries with RosettaLigand. PLoS One. 2015;10(7):e0132508.

115. Lyskov S, Chou FC, Conchuir SO, Der BS, Drew K, Kuroda D, et al. Serverification of molecular modeling applications: the Rosetta Online Server that Includes Everyone (ROSIE). PLoS One. 2013;8(5):e63906.

116. Kabuta T, Suzuki Y, Wada K. Degradation of amyotrophic lateral sclerosis-linked mutant Cu, Zn-superoxide dismutase proteins by macroautophagy and the proteasome. Journal of Biologial Chemistry. 2006;281(41):30524–33.

117. Wang Y, Mandelkow E. Degradation of tau protein by autophagy and proteasomal pathways. Biochemical Society Transactions. 2012;40(4):644–52.

118. Tobler AR, Constam DB, Schmitt-Graff A, Malipiero U, Schlapbach R, Fontana A. Cloning of the human puromycin-sensitive aminopeptidase and evidence for expression in neurons. J Neurochem. 1997;68(3):889–97.

